# Unconventional DNA architecture in a dopamine–bound aptamer complex

**DOI:** 10.1101/2025.11.27.690961

**Authors:** Emily Hoi Pui Chao, Eric Largy, Yunus A. Kaiyum, Minh-Dat Nguyen, Brune Vialet, Philippe Dauphin-Ducharme, Philip E. Johnson, Cameron D. Mackereth

## Abstract

Aptamers are oligonucleotides that have been selected to bind a particular target. Despite the growing popularity of functional DNA aptamers, there remains limited knowledge of their binding mechanisms as few have been characterized at the atomic level. Here we use NMR spectroscopy to obtain structural details of RKEC1, a shortened version of a DNA aptamer previously reported to bind dopamine. We find that RKEC1 forms a compact structure upon ligand binding that lacks any Watson-Crick duplex regions or G-quadruplex core, in stark contrast to nearly all predicted and observed DNA aptamer folds. The atomic details explain dopamine specificity amongst structurally similar compounds, and the determined DNA fold was used to guide biosensor design. The aptamer structure further suggests that DNA folding can access an extensive conformational landscape reminiscent of RNA, thus expanding the diversity traditionally captured by predictive algorithms.

## INTRODUCTION

Aptamers are single-stranded oligonucleotides that are selected from a large pool of random sequences to bind a target, which can be proteins, cells or small molecules.(Bock et al., 1992; DeRosa et al., 2023; Ellington & Szostak, 1990, 1992; Tuerk & Gold, 1990) The process by which aptamers are discovered is known as systematic evolution of ligands by exponential enrichment (SELEX). In initial SELEX experiments, the pool of oligonucleotides was passed over an immobilized target and the binding fraction subsequently amplified by PCR. More recently, a method called Capture-SELEX has been developed where instead the DNA library is immobilized and then exposed to the ligand, with the aptamer-ligand complex released upon binding the target.(Nutiu & Li, 2005; Yang et al., 2016) Aptamers discovered using Capture-SELEX often display large-scale folding with ligand binding, as a structural change accompanies the required displacement of the DNA upon ligand binding during selection. The resulting aptamers are often employed as recognition elements for diverse applications including the electrochemical aptamer-based (E-AB) sensing platform.(Huldin et al., 2025; Nguyen et al., 2025; Wu & Plaxco, 2025)

Using this approach, a recently developed DNA aptamer for dopamine (Nakatsuka et al., 2018) has been the focus of a number of recent studies, likely due to the importance of the target as a neurotransmitter. The aptamer was predicted to form a lower stem involving the nucleotides that bind the immobilization strand during the Capture-SELEX protocol, with a putative bulge and second stem in the variable region of the selected DNA. A later variant where a C-A mismatch in the stem was changed to a T-A Watson-Crick base pair (**Fig. 1a**, DA-mut3) was shown to modestly increase affinity.(Liu et al., 2021) This stabilized variant was also shown to improve the NMR spectra, with data indicative of a single structure formed upon binding dopamine in a 1:1 ratio.(Kaiyum et al., 2024) The NMR data also highlight a large amount of ligand-induced structure formation for both the long stem DA-mut3 and stem truncation variants.

**Fig. 1.**
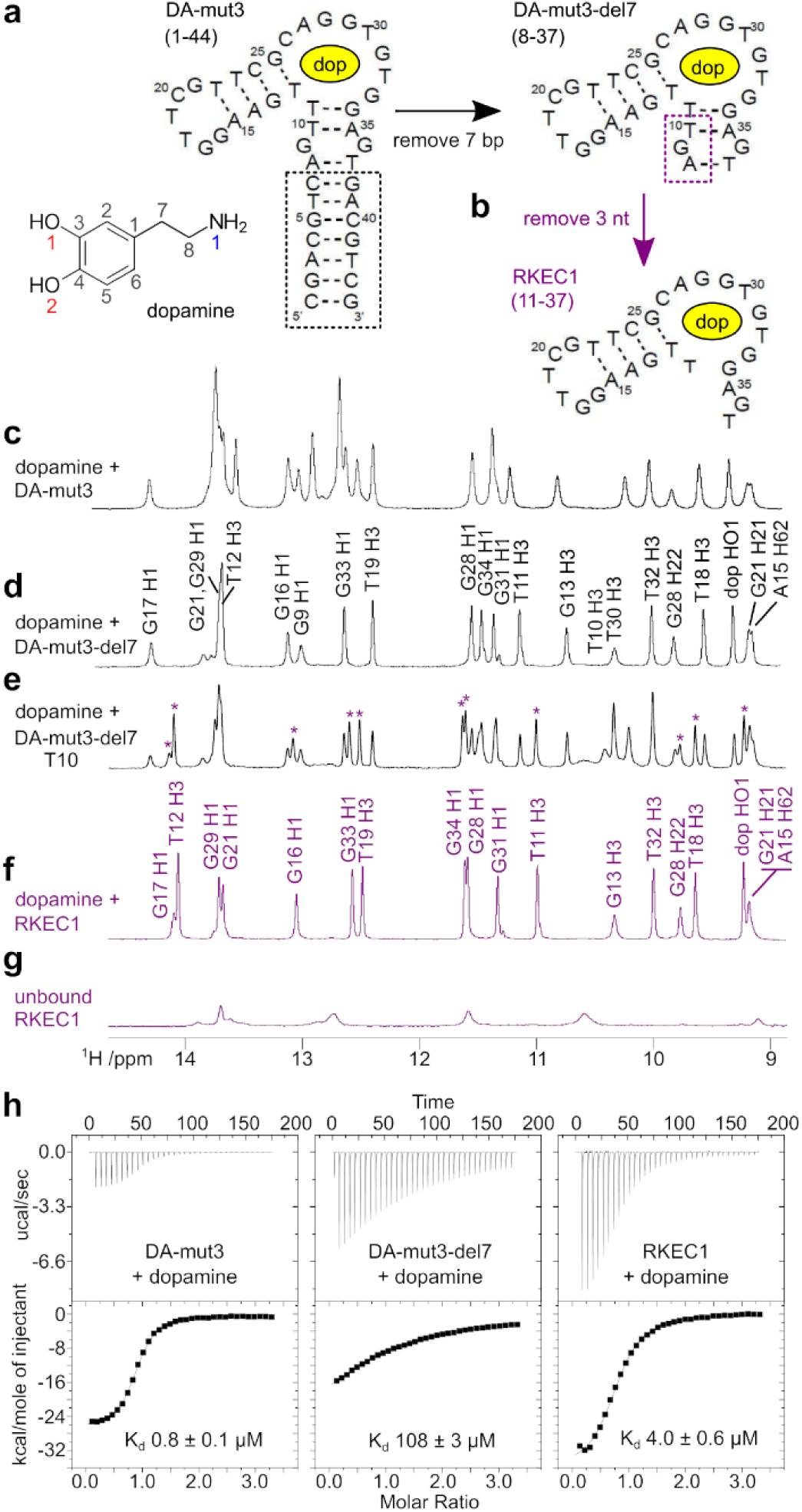
Aptamer truncation for structural study. (**a**) Putative secondary structures for DA-mut3 and truncated DA-mut3-del7 dopamine-binding DNA aptamers.(Kaiyum et al., 2024; Liu et al., 2021; Nakatsuka et al., 2018) (**b**) Further truncation of three 5′ nucleotides leads to the minimal dopamine-binding aptamer RKEC1. (**c**-**f**) The imino regions from 1D ^1^H NMR spectra of dopamine-bound aptamers depicted in (a,b). Additional NMR data and assignment information in **Supplementary Table 3**, **Supplementary Figs. 1-3**. (**g**) Imino region for the unbound RKEC1. (**h**) ITC thermograms and derived K_d_ values for dopamine binding to the aptamers depicted in (a,b). Additional thermodynamic data in **Supplementary Table 1**.

Despite advances, aptamers are often thought of as a “black box”. This is because little is known about how most aptamers function. Details of what structure they form and how the aptamer interacts and bind their target ligand are often unknown, and this is particularly true for DNA aptamers that target small molecules. To date, only a few DNA aptamers structures in complex with small molecules have been solved.(Andralojc et al., 2022; Lin & Patel, 1997; X. Lin et al., 2025; Liu et al., 2023; Robertson et al., 2000; Xu et al., 2023; Xu et al., 2022) Aside from the ochratoxin A-binding aptamer which presents a G-quadruplex motif, all others are stem-stabilized loop structures. While other structural motifs such as pseudoknots or three-way junctions have been suspected to provide a stable platform on which to build target recognition capabilities, those remain to be structurally resolved.

With the goal of understanding the atomic basis of aptamer recognition of dopamine, and to expand on the list of DNA aptamer structures, here we report the three-dimensional structure of dopamine-aptamer complex. We find that the DNA aptamer forms an unconventional compact structure in the presence of its ligand, with only two Watson-Crick base pairs formed while 11 non-Watson-Crick base pairs are present. The binding interactions seen in the dopamine-aptamer complex are consistent with solution-based biophysical characterization conducted on 12 structural analogues of dopamine, showcasing its intricate interaction with its target. The unusual fold of this DNA aptamer demonstrates the complexity of structures that can be attained by DNA aptamers to bind their targets, which we then use to accurately assign aptamer labelling sites for translation into the E-AB sensing platform.

For the purpose of structure characterization by NMR spectroscopy, we were interested in reducing the aptamer size to simplify the spectra while maintaining sufficiently high affinity. Using a series of truncations based on the 44-nt DA-mut3, we had previously observed that deletion of up to seven base pairs retained a similar binding mode for dopamine, as noted by comparing the ^1^H NMR spectra (**Fig. 1a**)(Kaiyum et al., 2024). This smaller dopamine-binding aptamer of 30 nt (DA-mut3-del7, **Fig. 1a**) binds dopamine with a reduced affinity of 108 µM (versus 0.8 µM for the parent DA-mut3; **Fig. 1e**, **Supplementary Table 1**), but with the advantage of fewer and better-separated peaks of the dopamine-bound aptamer (**Fig. 1d**). We therefore decided to use DA-mut3-del7 for structural studies and proceeded to assign the observed peaks. To simplify the assignment process and to avoid ambiguity, we chemically synthesized a series of oligonucleotides in which a single guanosine, thymidine or adenosine during synthesis was spiked with a uniform ^13^C,^15^N-labeled phosphoramidite (**Supplementary Table 2**). In an initial step, this strategy allowed us to assign all observable imino and amino ^1^H-^15^N cross peaks for the enriched nucleotides (**Supplementary Fig. 1**). At this stage, it was already evident that there were atypical DNA structures in the dopamine-aptamer complex, with few Gua H1 and Thy H3 resonances appearing in the usual duplex chemical shift range, and a surprisingly large number of observed amino resonances (**Fig. 1d**, **Supplementary Fig. 1**).

During the 3′ to 5′ chemical synthesis, we experienced some difficulty incorporating ^13^C,^15^N-labeled thymidine phosphoramidites, such that abortive DNA was produced at the nucleotide prior to the position chosen for labeling (**Supplementary Fig. 2**). These smaller oligonucleotides did not interfere with the assignment process as they did not form complexes with dopamine and were not enriched in ^13^C or ^15^N. Nevertheless, one of the 5′-truncated oligonucleotides did in fact form a stable complex, as observed with a second set of peaks in the 1D ^1^H NMR spectrum for the T10 sample (**Fig. 1e**). This DNA was missing the three 5′ nucleotides A8-G9-T10 (**Fig. 1b**) but was surprisingly able to form a complex with dopamine (**Fig. 1f**), despite our earlier truncation study revealing that removal of just the A8-T37 base pair significantly reduced dopamine binding (Kaiyum et al., 2024). Furthermore, we found that binding studies of this shortened aptamer (herein named RKEC1) corresponded to an unexpected increase in affinity as compared to DA-mut3-del7 (4 µM vs 108 µM, **Fig. 1e**) even though fewer nucleotides were present. As expected, the unbound RKEC1 did not exhibit any stable structure in the absence of the dopamine ligand (**Fig. 1g**). Due to the combination of improved affinity, further reduction in size, and an NMR spectrum consistent with similar dopamine recognition, we decided to switch to RKEC1 for structure determination. Near complete assignment of observable ^1^H signals was obtained by initial comparison to DA-mut3-del7 as well as additional NMR spectra and labeled samples (**Supplementary Table 3**, **Supplementary Fig. 3**).

### Overall structure of dopamine-bound RKEC1

Taking advantage of the NMR chemical shift assignments, we next used a two-part strategy to determine an ensemble of solution structures for dopamine-bound RKEC1. The first step relied on an iterative NOE assignment and structure calculation procedure with ARIA2.3/CNS1.2 (Brunger, 2007; Rieping et al., 2007) using NOESY spectra acquired at 298 K and 278 K, along with dihedral restraints. The structure statistics from ARIA/CNS are shown in **Table 1**, with the 43 ligand-DNA intermolecular restraints detailed in **Supplementary Table 4**. In the second step, the 15 models were each subjected to 10 ns of restrained molecular dynamics (**Supplementary Fig. 4**) in AMBER24(Case et al., 2023) using explicit water and buffer ions as described in the Methods. The final structure statistics are presented in **Table 1**, along with visualization of distance restraints (**Supplementary Fig. 5**), pairwise analysis (**Supplementary Fig. 6**) and sugar pucker distribution (**Supplementary Fig. 7**).

**Table 1.** NMR and refinement statistics for the ensemble of RKEC1 bound to dopamine. . Structure determination using ARIA/CNS followed by AMBER. Statistics calculated over all 15 models.

| **ARIA/CNS** |  |  |
| --- | --- | --- |
| Distance restraints |  |  |
| Total NOE | 963 |  |
| Intra-residue | 502 |  |
| Sequential ($\vert i - j\vert = 1$) | 184 | |
| Nonsequential ($\vert i - j\vert > 1$) | 144 | |
| Intermolecular | 43 |  |
| Ambiguous <sup>a</sup> | 90 |  |
| Dihedral angle restraints |  |  |
| $\chi^1$ | 27 | |
| Sugar pucker | 16 |  |
| **Structure statistics** |  |  |
| Violations (mean and s.d.) |  |  |
| Distance constraints (Å) | 0.009 ± 0.002 |  |
| Dihedral angle restraints (°) | 0.29 ± 0.05 |  |
| Deviations from idealized geometry |  |  |
| Bond lengths (Å) | 0.002 ± 0.000 |  |
| Bond angles (°) | 0.37 ± 0.01 |  |
| Impropers (°) | 0.202 ± 0.003 |  |
| Average pairwise r.m.s. deviation |  |  |
| All RNA heavy (Å) | 0.8 ± 0.1 |  |
| **AMBER <sup>b</sup>** |  |  |
| **Structure statistics** |  |  |
| Violations (mean and s.d.) |  |  |
| Number of distance violations (> 0.3 Å) | 0 |  |
| Average pairwise r.m.s. deviation |  |  |
| DNA backbone heavy atoms (Å) | 1.3 ± 0.3 |  |
| All heavy atoms (Å) | 1.3 ± 0.3 |  |

<sup>a</sup> Class of distance restraints in Aria2.3 for cross peak volumes contributed by two or more overlapping NOE
<sup>b</sup> Only the 873 non-ambiguous distance restraints were used for the AMBER minimization protocol, with no dihedral restraints

The ensemble of 15 calculated structures of dopamine-bound RKEC1 (**Fig. 2a**) has an all-heavy atom r.m.s. deviation of 1.3 ± 0.3 Å. Most notable was the unusual DNA fold within the complex, as illustrated with a representative model closest to the average structure (**Fig. 2b**) and a simplified schematic (**Fig. 2c**). In a dramatic contrast to the predicted DA-mut3 secondary structure (**Fig. 1a**), the dopamine-binding aptamer does not display regions of duplex DNA. Instead, the folding arrangement is such that the 5’ end is located midway along the structure as it forms part of a staggered triplex element instead of the predicted duplex stem. Another significant feature is a loop region that folds back on itself, anchored by a Watson-Crick base pair between C26 and G33, that creates an additional region of three-and four-base planes. The dopamine ligand lies on top of the C26-G33 base pair between these two large regions and is enclosed by the folded-back loop, with stacking of the dopamine aromatic ring between bases G33 and T32 (**Fig. 2d**). The dopamine hydroxyl groups form hydrogen bonds with oxygen atoms and an imino hydrogen from the T18 and T19 bases, with additional ionic interactions between the dopamine amino group and nucleotides A27 and G28.

**Fig. 2.**
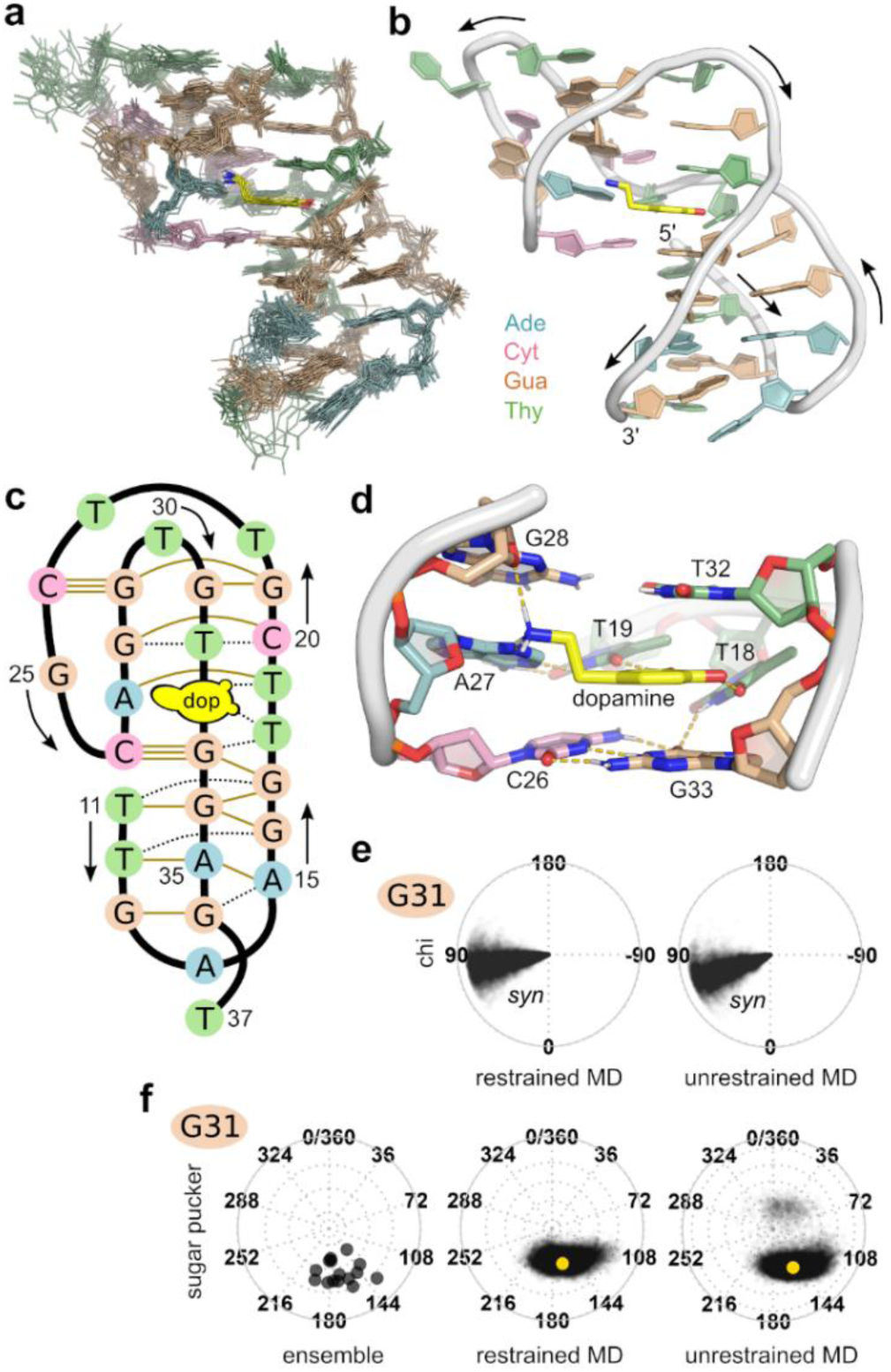
Structure of dopamine-bound RKEC1. (**a**) Ensemble of 15 dopamine-bound RKEC1 solution structures determined using ARIA1.2/CNS2.3 followed by AMBER24. See Methods for complete details and Table 1 for structure statistics. (**b**) Representative closest-to-average model shown as a cartoon, with arrows to indicate direction of 5′ to 3′. (**c**) Schematic of dopamine-bound RKEC1 with base information, residue number annotation, and direction of 5′ to 3′ indicated. Gold lines indicate major base pairs, with dotted black lines showing base pairs connected by a single hydrogen bond or not present in all models. The two Watson-Crick C–G base pairs are indicated with three connecting lines. (**d**) Close-up view of the dopamine binding site in RKEC1. Dotted lines indicate hydrogen bonds. (**e**) Chi dihedral angle values for G31 extracted from 1-μs restrained and unrestrained molecular dynamics simulations in AMBER 24. The center position indicates the start of the simulation which then proceeds outwards to the border. Full analysis in **Supplementary Figs. 9, 12**. (**f**) Sugar pucker observed for G31 extracted from the ensemble of 15 structures (**Supplementary Fig. 7**), as well as 1-μs restrained and unrestrained molecular dynamics simulations in AMBER 24 (**Supplementary Figs. 9,12**). From center to border represents increasing amplitude of the sugar pucker, with planarity in the center.

Comparison of the 15 models in the ensemble indicates only a limited degree of variation in the structural details (**Fig. 2a**). To further probe the potential dynamics of the complex, we ran four separate 1-µs MD simulations in AMBER with the NMR distance restraints active (**Supplementary Fig. 8a**). Analysis of the separate trajectories revealed similar behavior for dihedral angles, sugar pucker, and the formation of key hydrogen bonds (**Supplementary Figs. 9-11**). Using G31 as an example, as this guanine is the only RKEC1 base in the *syn* conformation (**Supplementary Fig. 3b)**, this dihedral angle remains stable throughout the restrained MD simulations (**Fig. 2e**). Similarly, the G31 sugar pucker values observed in the restrained MD simulations align with the range from the calculated ensemble (**Fig. 2f**). An additional 1–µs simulation performed in the absence of any restraint resulted in comparable structure parameters over the trajectory (**Fig. 2e,f**, **Supplementary Figs. 12-14**), highlighting the stable nature of the DNA fold and DNA-ligand interactions. Also observed in each of the four simulation replicates was the relatively stable position of a sodium ion that may partially stabilize the loop nucleotides C24, G25, A27, G28, and G29 (**Supplementary Fig. 15**). If present in solution, this ion would likely represent a magnesium ion from the buffer and could explain the increased affinity of the aptamer for dopamine in the presence of MgCl_2_. (Liu et al., 2021; Nakatsuka et al., 2021)

### Abundance of atypical DNA base interactions

The most notable feature of the dopamine-bound RKEC1 DNA is the lack of canonical base-pair interactions. Surprisingly, there are only two Watson–Crick base pairs in the entire complex: the C26–G33 base pair that stabilizes the loop position, and a C24–G29 base pair located further within the loop. To help characterize the remaining base pairs, we therefore turned to the nomenclature strategies developed for RNA structure analysis for which a full range of base-pair types can be described. Using the program DSSR(Lu et al., 2015) we identified 16 base pairs that appear in all 15 models (**Supplementary Table 5**). Thirteen of these base pairs are connected by at least two hydrogen bonds, and as such they can be annotated using the Saenger base-pair nomenclature created for RNA.(Saenger, 1984) Using the Saenger nomenclature to annotate the RKEC1 DNA (**Fig. 3a**), it is possible to describe how these noncanonical base pairs contribute to the overall DNA architecture within the complex.

**Fig. 3.**
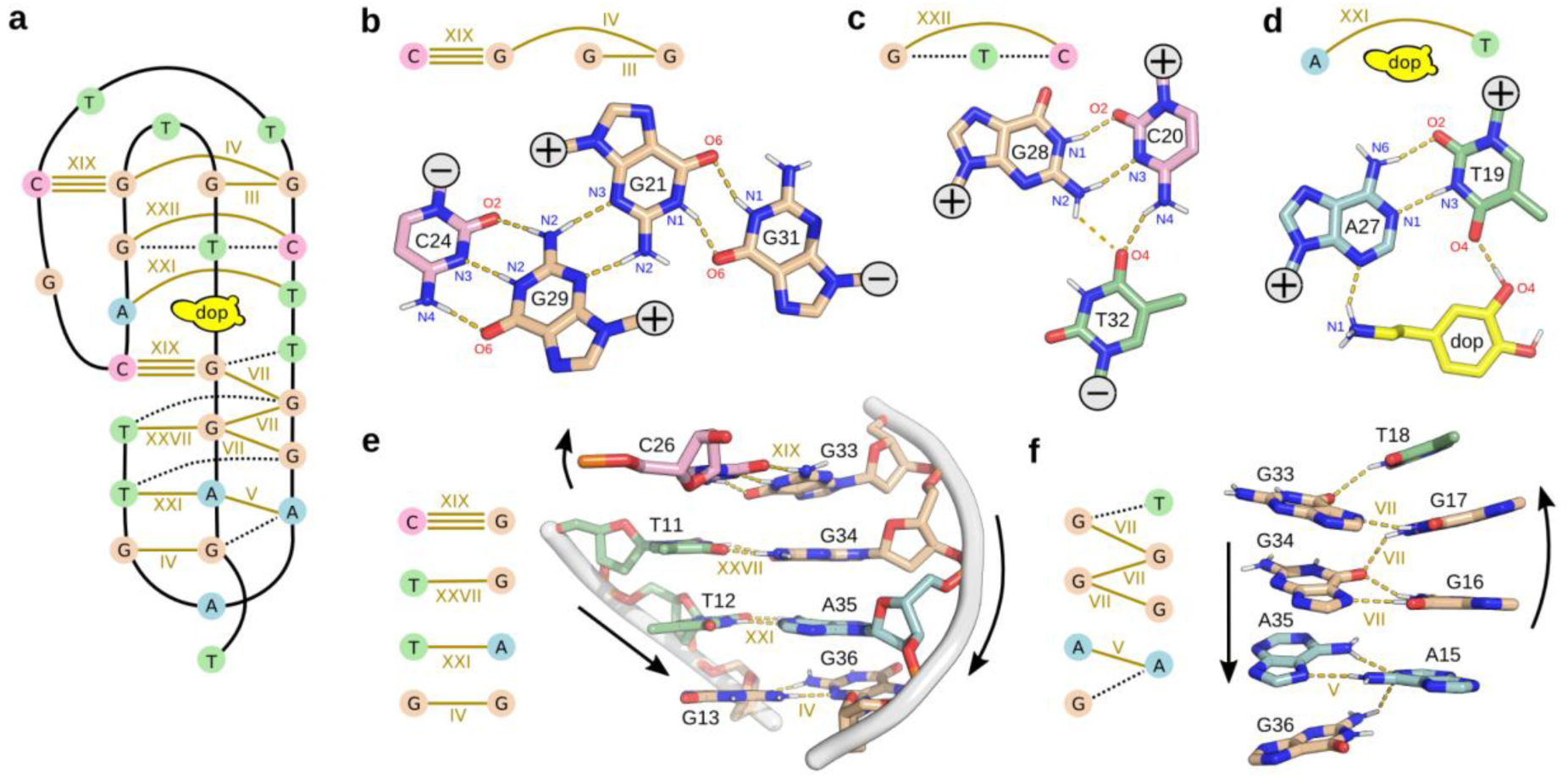
Diverse range of DNA base-pairs observed in dopamine-bound RKEC1. (**a**) Schematic of dopamine-bound RKEC1 as in Fig. 2c, but with the major base pairs annotated using the Saenger nomenclature.(Saenger, 1984) Additional details and nomenclature based on the RNA Leontis-Westhof(Lemieux & Major, 2002; Leontis & Westhof, 2001) and DSSR(Lu et al., 2015) systems are included in **Supplementary Table 5**. (**b-d**) Close-up views of three base-pair regions taken from (**a**), with annotation of the hydrogen-bond heavy atoms. The 5′ to 3′ direction of the DNA strand for each nucleotide is indicated in the gray circle (+, out of page;-, into page). (**e**) Close-up view of the four stacked base-pairs located below the bound dopamine. The 5′ to 3′ direction of the DNA strand is indicated for each segment. (**f**) Close-up view of the staggered base pair region, annotated as in (**e**)

Starting from the top of the complex (**Fig. 3b**), a four-base plane is formed by the antiparallel C24–G29 Watson–Crick (Saenger nomenclature XIX) and G21–G31 symmetric N1-carbonyl (III) pairs, connected by a parallel G21–G29 symmetric N3-amino (IV) base pair. The next plane is a base triple (**Fig. 3c**), primarily formed by the reverse Watson–Crick (XXII) C20–G28. A second reverse Watson–Crick base pair (T19–A27) forms part of a pseudo base-triple that incorporates the bound dopamine (**Fig. 3d**). The remaining base pairs stabilize the staggered base-triple region. In detail, the first three bases of RKEC1 form a series of parallel base pairs (**Fig. 3e**) that include T11–G34 (XXVII, reverse wobble), T12–A35 (XXI, reverse Watson–Crick) and G13–G36 (IV, symmetric N3-amino). A fourth base pair that completes this segment includes the isolated C26 base from the loop region, allowing for an antiparallel conformation to form the C26–G33 Watson–Crick base pair. Finally, four bases from A15 to T18 contact the three bases from G33 to A35 in an antiparallel but staggered orientation (**Fig. 3f**), starting with a N1-amino, N7-amino (V) A15–A35 base pair followed by three G–G base pairs with N1-carbonyl, N7-amino orientation (VII). In addition, there are at least six recurring base pairs identified in the ensemble that are connected by single hydrogen bonds to further stabilize the RKEC1 fold (**Supplementary Table 5**, and dotted lines in **Fig. 3a,c,f**).

### Molecular basis of dopamine specificity

In addition to providing details of the DNA fold, the structure of the dopamine-bound RKEC1 allows for the characterization of the ligand-binding mechanism. The dopamine molecule is recognized by several features as summarized by LigPlot+ (**Fig. 4a**).(Laskowski and Swindells 2011) The aromatic ring of dopamine stacks between the bases T32 and G33, with additional hydrophobic contacts primarily between dopamine H2 and A27 H2. Hydrogen bonds connect O4 from T18 with both dopamine hydroxyls, with an additional hydrogen bond between T19 O4 and dopamine O1. Finally, the dopamine amino group forms numerous ionic interactions with base and sugar atoms of A27 and G28. Upon binding to dopamine, the ligand is mostly buried within the complex, with only H6, H71/H72 and H81/H82 exposed to the solvent (**Fig. 4b**).

**Fig. 4.**
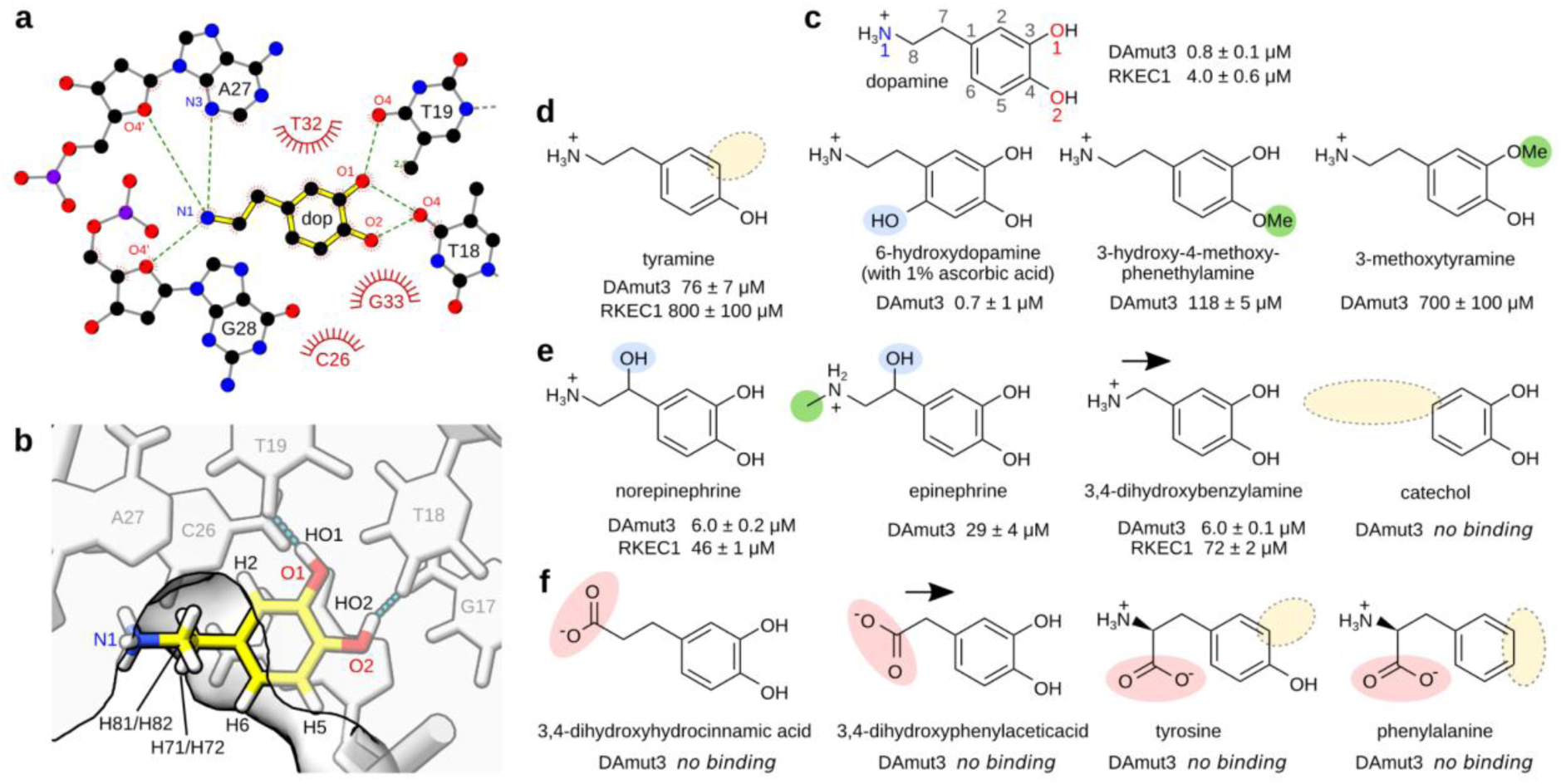
Probing the dopamine binding site with ligand analogs. (**a**) Dopamine-aptamer interactions identified with LigPlot+,(Laskowski & Swindells, 2011) including hydrogen bonds (dotted lines and heavy atom annotation) and non-bonded contacts (spoked arcs). (**b**) Cross-section surface representation of RKEC1 illustrates varying degrees of solvent accessibility for atoms in the bound dopamine. (**c**) ITC reference affinity of dopamine for the RKEC1 aptamer and the parent DA-mut3 (Fig. 1e, **Supplementary Table 1**). Dopamine heavy atoms are annotated by number. (**d**) Effect of the removal or addition of hydroxyl and methoxy groups around the dopamine aromatic ring emphasize the role of key hydrogen bonds. (**e**) Modification of the aliphatic segment has moderate effects on affinity, but complete removal of the aliphatic amine eliminates binding. (**f**) No binding is observed when the amine group is replaced by a carboxylate, even with retracting the group away from the phosphate backbone. No binding was also observed for the amino acids tyrosine or phenylalanine. Complete binding parameters for all analogs in **Supplementary Table 1**, with representative thermograms in **Supplementary Fig. 16**.

Based on this atomic information, we decided to probe the specificity of ligand binding by measuring the affinity of a series of dopamine analogues by using ITC (**Supplementary Table 1**, **Fig. 4c**, **Supplementary Fig. 16**). DA-mut3 was included in this analysis as it fully preserves the binding pocket to RKEC1 based on NMR analysis, but has a higher intrinsic affinity to allow for the characterization of weaker ligands. Our first series of analogues probed the importance of the dopamine hydroxyl groups (**Fig. 4d**). The dopamine hydroxyl at O1 participates in hydrogen bonds to T18 O4 and T19 O4 (**Fig. 4a**). Removal of this hydroxyl in tyramine significantly reduces affinity to DA-mut3 by a factor of 100 (76 µM) and by a factor of 200 for RKEC1 (800 µM), highlighting the importance of these hydrogen bonds in ligand binding. In contrast, addition of a new hydroxyl on C6 in 6-hydroxydopamine does not affect affinity, in keeping with the fact that this position would be solvent exposed (**Fig. 4b**). Adding a methyl group to either hydroxyl group would also be predicted to perturb hydrogen-bond formation and add steric penalties, and accordingly 3-hydroxy-4-methoxy-phenethylamine (methyl ether at O2) reduces binding by a factor of 150 (118 µM) and 3-methoxytyramine (methyl ether at O1) strongly reduces binding by almost three magnitudes (700 µM).

The aliphatic amino moiety was also evaluated for its importance in ligand binding (**Fig. 4e**). Given the exposure of H71/H72 to the solvent (**Fig. 4b**), addition of an hydroxyl group at C7 was not expected to significantly alter binding, and indeed norepinephrine binds with only a modest reduction in affinity to DA-mut3 (6 µM) or RKEC1 (46 µM). The close biological analogue epinephrine has an additional amino-methylation which further reduces affinity (29 µM for DA-mut3) likely due to perturbation of ionic contacts between the amino group and nucleotides A27 and G28 (**Fig. 2d**, **Fig. 4a**). The removal of one methylene group in 3,4-dihydroxybenzylamine modestly reduces affinity, illustrating that the position of the amino group within the RKEC1 complex is also important. The complete removal of the aliphatic amine in catechol prevents any binding, supporting a key role for this moiety in DNA-ligand interactions. The importance of the dopamine terminal amino group is also illustrated by a final series of analogues in which the amino group is replaced by a carboxyl group (**Fig. 4f**). For 3,4-dihydroxyhydrocinnamic acid and 3,4-dihydroxyphynylacetic acid, which directly compare to dopamine and 3,4-dihydroxybenzylamine, the presence of the negative charge completely prevents binding even though the remainder of the molecule remains intact. Finally, the two amino acids tyrosine and phenylalanine partially resemble dopamine and retain the amino group, but the addition of a terminal carboxyl group and lack of one or both aromatic hydroxyls also abolished binding to the DA-mut3 aptamer.

### Structure-guided design of an E-AB biosensor

The structure of the dopamine-bound RKEC1 aptamer was unexpected based on secondary structure predictions, and we decided to take advantage of this atomic information to guide design of the aptamer sequence for biosensing. The common strategy to convert an aptamer into an electrochemical aptamer-based (E-AB) biosensor is to attach a thiol anchor at the 5′ end and the methylene blue (MB) redox reporter at the 3′ end (**Fig. 5a**, **Supplementary Table 6**).(Arroyo-Currás et al., 2017; Downs & Plaxco, 2022; Wu et al., 2023).

**Fig. 5.**
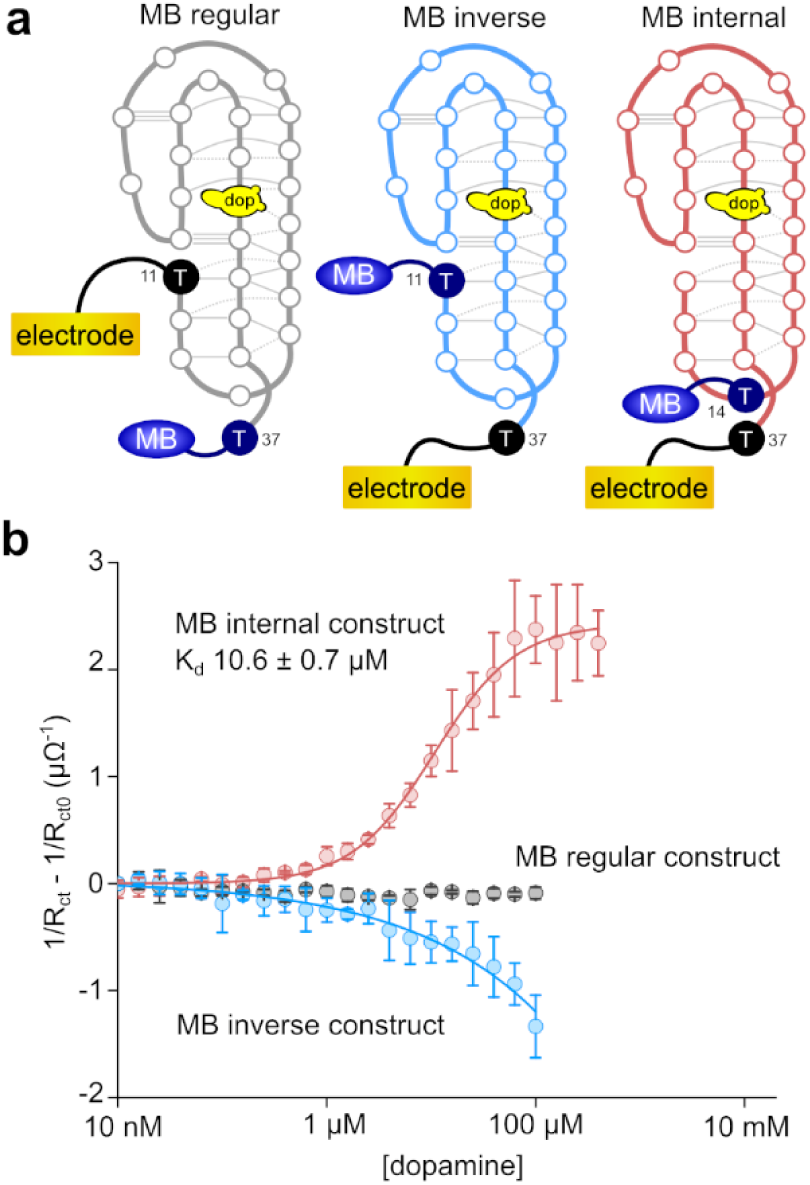
**Electrochemical aptamer-based (E-AB) biosensors based on RKEC1**. (**a**) Three different biosensor designs with variation in the position of the methylene blue redox reporter (MB) and location of the thiol anchor to the gold electrode. RKEC1 is expected to be disordered or exhibit conformational heterogeneity in the absence of dopamine. Oligonucleotide sequence details in **Supplementary Table 6**. (**b**) E-AB sensor responses to increasing dopamine concentrations using the three RKEC1 sensor designs showed either increase or decrease in measured charge transfer resistances associated with changes in the distance or environment of the redox reporter, as seen in the corresponding phase angle shift response (**Supplementary Fig. 17**). Only the MB internal variant produced a ligand-dependent response. Measurements performed in 1X PBS buffer with 2 mM MgCl_2_ as well as 0.02 % ascorbic acid to prevent dopamine oxidation. Error bars illustrate the standard deviation of signals produced by three sensors independently fabricated and tested.

Using this regular arrangement, dopamine binding was interrogated using electrochemical impedance spectroscopy (see Methods for details).(Downs et al., 2020; Nguyen et al., 2024) This arrangement however did not result in any dopamine-dependent biosensor response (**Fig.5b**, MB regular construct). A simple reversal of the thiol anchor and redox reporter, previously found to increase E-AB responses (Chamorro-Garcia et al., 2021), only modestly improved the response to dopamine (**Fig.5b**, MB inverse construct). By inspecting the NMR data and structure of the RKEC1 complex, we noted that the unbound RKEC1 configuration is predominantly disordered and the interaction with dopamine is coupled with structure formation. Our two E-AB designs indeed took advantage of the initial disordered state, and therefore the problem likely pertained to the inability of the modified aptamer to bind dopamine, or a dopamine-binding region distal to the gold surface.

An analysis of T11 at the 5′ end of RKEC1 indicates that part of the affinity improvement over DA-mut3-del7 might be due to the 5′ hydroxyl group at the 5′ end in place of a phosphate or additional nucleotides (**Supplementary Fig. 17a**). A phosphate in this position would bring a destabilizing negative charge in close proximity to the phosphates around T19. It would also suggest that appending anything to the 5′ end could similarly disrupt the ability of RKEC1 to form a stable folded complex. This would explain why attempts to anchor the aptamer at the 5′ end in the regular construct abolished all sensor response (**Fig. 5b**). The RKEC1 structure also indicates that although the methylene blue may be more tolerated at the 5′ end, it would not be near the sensor surface in the ligand-bound conformation (**Fig. 5a**). Fortunately, analysis of the structure suggested that A14 would be closer to the sensor surface in the bound conformation, and importantly this adenine did not make specific interactions within the complex. NMR analysis confirms that changing this base (A14G) does not affect ligand binding (**Supplementary Fig. 17b**).

Using this atomic information, we designed a third arrangement that incorporates the redox reporter internally at the A14 position (swapping for a thymine-modified methylene blue) with the thiol anchor at the 3′ end (**Fig. 5a**). The thiol anchor would now be in the vicinity of the redox reporter to allow for a shorter reporter-surface distance and increase the ability to resolve changes in electron transfer (Nguyen et al., 2025). This new, and perhaps atypical, internally-modified RKEC1 design successfully produced a concentration-dependent dopamine response with a *K_d_* value of 10.6 ± 0.7 µM (**Fig. 5b**). This measured affinity was similar to that of the unmodified aptamer (4 µM). We attributed this dose-dependent signal (**Supplementary Fig. 17c**) to the transition of the redox reporter from a more distal position to within the proximity of the electrode upon binding dopamine because the unbound state of the aptamer is largely disordered.

## DISCUSSION

Here we report on the structure of the DNA aptamer RKEC1 in complex with dopamine. When bound, RKEC1 forms a compact structure where non-Watson–Crick base-pair formation predominates. In the structure, only two Watson–Crick base pairs form. In contrast, we identified 11 non-Watson–Crick base pairs involving a variety of base combinations, along with additional pairings that involve a single hydrogen bond (**Supplementary Table 5**). Furthermore, the two Watson–Crick base pairs are not sequential, meaning that there is no region of standard duplex in the folded DNA. Although perhaps an extreme case, the RKEC1 structure illustrates a complexity in DNA conformation that is not currently addressed in predictive tools for DNA secondary and tertiary structures.

In the recent Critical Assessment of Techniques to Protein Structure Prediction (CASP16), we had included the RKEC1–dopamine complex as target D1273 within both the nucleic acid and ligand categories.(Kretsch et al.) Whereas the overall prediction accuracy was high for the RNA targets, the lack of any accurate predictions for RKEC1 (the only DNA target) was notable. One difficulty in the RKEC1 prediction may be the prevalence of non-standard base pairing that is currently missing for DNA structures deposited in the PDB. This situation is however more common in RNA structures (Holbrook, 2005; Vicens & Kieft, 2022) and some of the predictions did have limited regions of non-standard base pairs. As part of the modelling process, the currently small dataset of complex DNA structures hinders proper training strategies, and this problem can only be improved with more experimental structures. In addition, numerous DNA–ligand interactions are present within the RKEC1 complex, and this may also present difficulties when predicting the DNA fold in the absence of dopamine.

Related to this last point, the dopamine DNA aptamer is one of only a few aptamers selected using Capture-SELEX that have had their structures determined, the others being the theophylline aptamer (Xiaowei Lin et al., 2025) and an ATP-binding aptamer (Jiang et al., 2025). In capture-SELEX it is the DNA library and not the ligand that is immobilized. It would therefore be expected that the ligand is more enveloped or buried in the aptamer structure than seen in earlier reported structures of DNA aptamers derived from other SELEX methods. This property is indeed seen in the RKEC1–dopamine complex where extensive hydrogen bonding and stacking interactions between the ligand and aptamer are seen (**Fig. 4b**), with only a small portion of the dopamine solvent exposed. The burial of the ligand within the aptamer is likely related to the ligand-induced folding seen with this aptamer where the free aptamer shows only a few weak imino signals (**Fig. 1g**), requiring ligand addition to trigger aptamer folding (**Fig. 1f**).

The small size, presence of hydrogen-bond donors and acceptors, and the aromatic nature of dopamine make it a good candidate to participate in DNA interactions that partially mimic nucleobase interactions. One such binding mechanism involves the stacking of dopamine between bases T32 and G33 (**Fig. 2d**) and another mechanism involves contribution of dopamine to form a pseudo base triple with the T19–A27 base pair (**Fig. 3d**). These two properties are also observed in the DNA aptamer recognition of theophylline and related molecules.(X. Lin et al., 2025) The binding pocket for this series of ligands involves stacking between a guanine and cytosine base, with hydrogen bonds to an adenine and thymine in a planar arrangement. This work further illustrates the use of structural details to create aptamers with altered binding specificity.

We also demonstrate here that structural knowledge can be critical for designing aptamer-based biosensors. E-AB biosensors are a common use of aptamers in biosensor devices and typically involves the immobilization of the aptamer on a gold surface with a thiol group attached to the 5’end of the DNA and a redox reporter, methylene blue, attached to the 3’ end. When this arrangement was used for the dopamine aptamer the resulting sensor did not work. From examining the structure, knowing that the 5’ end was not exposed while the 3’ end was more exposed, the location of the attachment was swapped resulting in a moderately functional sensor (**Fig. 5**). This sensor was then improved by noting that A14 is in a loop region, and not part of the structured core of the aptamer, and this residue should be close to the surface when immobilized by a thiol at the 3’ end of the aptamer. This adenosine was replaced with a thjymine to which the MB was attached and the resulting sensor worked efficiently (**Fig. 5**). This improved sensor demonstrated that a structure-guided approach is an effective approach to biosensor design.

In summary, we have determined the structure of a dopamine-binding DNA aptamer demonstrating that this aptamer has a complex architecture where non-Watson–Crick base pairs predominate. The aptamer envelops the ligand making extensive hydrogen bonds and stacking contacts. We also demonstrated the utility of a structure-guided approach to biosensor design.

## METHODS

### Oligonucleotides

Unmodified DNA oligonucleotides were purchased from Genscript and Integrated DNA Technologies. The series of site-specific [5 %-^13^C,^15^N]-enriched oligonucleotides (detailed in **Supplementary Table 2**) were synthesized at a scale of one micromole on an H-8 automated DNA synthesizer (K&A Labs, Germany). Starting with 1000 Å controlled pore glass solid support (SynBase CPG, Link Technologies, UK), the nucleotides were incorporated using the classic β-phosphoramidite methodology. Standard monomers (Bz-dA, dT, dmf-dG and Ac-dC) and solvents were purchased from Glen Research (USA). The uniformly-labeled (U-^13^C, U-^15^N, CP 95 %) dmf-dG, and dT phosphoramidites were purchased from Silantes (Germany). The deprotection and cleavage of the oligonucleotides were performed by incubation in ammonium hydroxide at 55 °C for 4h. Following an overnight removal of liquid by using a SpeedVac vacuum concentrator, the oligonucleotides were resuspended in 500 μL water. The samples were then exchanged into a buffer containing 20 mM sodium phosphate (pH 7.4), 140 mM NaCl and 2 mM MgCl_2_, by using a NAP-5 column (Cytiva). The DNA concentrations were determined by absorbance at 260 nm, with extinction coefficients provided by the OligoAnalyzer program (Integrated DNA Technologies).

### Isothermal titration calorimetry

Aptamer samples for ITC analysis were obtained from Integrated DNA Technologies (IDT, Coralville, Iowa) with standard desalting and used without further purification. The identity of the DNA was verified by mass spectrometry performed by the manufacturer. DNA samples were dissolved in distilled, deionized water and then exchanged three times in a 3 kDa molecular weight cutoff concentrator with 1 M NaCl and washed at least three times with distilled deionized water. All DNA samples were exchanged with the buffer (20 mM sodium phosphate, 140 mM NaCl, 2 mM MgCl_2_, pH 7.4) three times before use. Aptamer concentrations were determined by absorbance spectroscopy using the extinction coefficients provided by IDT. All small molecule ligands were obtained from Sigma Aldrich. ITC binding experiments were performed using a MicroCal VP-ITC instrument in a manner described previously.(Slavkovic et al., 2018; Slavkovic & Johnson, 2023) In brief, samples were degassed before analysis with a MicroCal ThermoVac unit. All experiments were corrected for the heat of dilution of the titrant. Titrations were performed with the aptamer samples in the cell, and the ligand as the titrant, in the needle. All aptamer samples were heated in a 95 °C water bath for 3 min and cooled in an ice water bath prior to use in a binding experiment to allow the DNA aptamer to anneal in an intramolecular fashion.

The binding experiments were performed at 20 °C with the aptamer solution at 20–60 μM aptamer concentration using a ligand concentration of 312 to 936 μM. Ligand solutions were diluted to experimental concentrations with buffer A. All binding experiments were performed at 20 °C and consisted of an initial delay of 60 s, first injection of 2 μL and 300 s delay. The subsequent 34 injections were 8 μL, spaced every 300 s to allow for adequate mixing and equilibration before the next injection. The first point was removed from all datasets due to the different injection volume and delay parameters. ITC data were fit to a one-site binding model using the manufacturer provided Origin 7 software. Representative thermograms appear in **Fig. 1h** and **Supplementary Fig. 16**.

### NMR spectroscopy

NMR data were collected on a Bruker Avance Neo 700 or 800 MHz spectrometer, equipped with triple-resonance gradient room-temperature or cryogenic probe, respectively. NMR data were processed by using NMRPipe/NMRDraw software (Delaglio et al., 1995) and NMR spectra were analyzed using Sparky (T. D. Goddard & D. G. Kneller, University of California, San Francisco, USA). Samples were measured at 278 K and 298 K, typically prepared in 20 mM sodium phosphate (pH 7.4), 140 mM NaCl and 2 mM MgCl_2_, with 10 % (v/v) D_2_O added for the lock. To prepare the samples, 1.2 molar equivalents of freshly prepared 100 mM dopamine in the above NMR buffer was added to each purified oligonucleotide sample. The samples were heated for 2 min at 95 °C then cooled rapidly in ice water. When necessary, the concentration of the aptamer-ligand complex was increased by using Amicon 3 kDa concentrators, with 170 μL of each final sample transferred to a 3 mm NMR tube. An initial 1D ^1^H spectrum was acquired for each sample, using excitation sculpting to remove the water signal, to ensure that complexes were correctly formed.

### Chemical shift assignment

Assignment of the dopamine-bound DNA aptamer was first determined for the DA-mut3-del7 oligonucleotide in complex with dopamine. The samples were prepared in 20 mM sodium phosphate (pH 7.4), 140 mM NaCl, 2 mM MgCl_2_ using 90% H_2_O/10% D_2_O or 100% D_2_O, with data acquired at 278 K and a field strength of 800 MHz. An initial 1D ^1^H NMR spectrum (Bruker pulse program *zgesgp*) was collected on a sample of 1.5 mM DA-mut3-del7 and 1.75 mM dopamine in 90% H_2_O/10% D_2_O to confirm complex formation, followed by 2D DQF-COSY (*cosydfgpph19*) and 2D ^1^H,^1^H-TOCSY (*dipsi2gpph19*, 80 ms mixing time) spectra. For better resolution of the sugar resonances to help in assignment, 2D 1H-1H NOESY (*noesyesgpph*, 200 ms mixing time) and 2D ^1^H,^1^H-TOCSY (*dipsi2gpph19*, 80 ms mixing time) spectra were acquired on a sample of 1 mM DA-mut3-del7 with a large excess of dopamine (2 mM) prepared in 100% D_2_O. Early in the data analysis it was evident that dopamine-bound DA-mut3-del7 lacked typical duplex signatures, and therefore to prevent any errors in assignment we generated a series of oligonucleotides in which a single nucleotide was enriched with uniform 5% [^15^N-^13^C]-labelling. Starting with guanosine labelling, we made 90% H_2_O/10% D_2_O samples of 2 mM DA-mut3-del7 and 2.2 mM dopamine in which G9, G13, G16, G17, G21, G25, G28, G29, G31, G33, G34 or G36 were separately enriched with ^13^C and ^15^N. This enrichment allowed us to acquire sensitivity-enhanced ^1^H,^15^N-HSQC with a ^15^N spectral width of 2500 Hz, centered on 150 ppm or 75 ppm, for detection of guanine base H1-N1 imine or H21/H22-N2 amine cross peaks, respectively. ^1^H and ^13^C sugar resonances for the labelled nucleotides were partially assigned based on 2D ^1^H,^13^C-HSQC spectra (*hsqcetgpsisp2.2*, with a ^13^C spectral width of 30 ppm centered on 75 ppm). In a similar approach with thymidine labelling, 90% H_2_O/10% D_2_O samples of 2 mM DA-mut3-del7 and 2.2 mM dopamine were prepared in which T10, T11, T12, T18, T19, T22, T23, T30 or T32 were separately enriched with ^13^C and ^15^N. ^1^H,^15^N spectra for the assignment of thymine base H3-N3 imines used a ^15^N offset of 160 ppm, with no change to the ^1^H,^13^C-HSQC parameters. A final adenosine series used separate enrichment of A8, A14, A15, A27 or A35 with the collection of ^1^H,^13^C-HSQC spectra, and included ^1^H,^15^N spectra centered on 75 ppm in order to detect possible H61/H62-N6 amine cross peaks. To assign possible cytosine base H41/H42-N4 amine cross peaks, a natural abundance ^1^H,^15^N-HSQC spectrum was collected on the T23 sample since it lacked an observable H3-N3 cross peak, but now with ^15^N centered on 90 ppm. These site-specific ^1^H-^15^N assignments are illustrated in **Supplementary Fig. 1**. Upon identification of RKEC1 as having higher dopamine-binding affinity than DA-mut3-del7, the assignment process was switched to dopamine-bound RKEC1. The partial chemical shift assignments for dopamine-bound DA-mut3-del7 were deposited in the BMRB under accession code 52688.

Due to the similar NMR spectra observed for dopamine-bound DA-mut3-del7 compared to the RKEC1 complex, it was possible to directly transfer most of the already determined chemical shift assignments. For spectra collected at 278 K, this was first accomplished with a natural abundance sample of 2 mM RKEC1 with 2.25 mM dopamine, prepared in 20 mM sodium phosphate (pH 7.4), 140 mM NaCl, 2 mM MgCl_2_ using 90% H_2_O/10% D_2_O. A pair of sensitivity-enhanced ^1^H,^15^N-HSQC with a ^15^N spectral width of 2500 Hz, centered on 150 ppm or 75 ppm, allowed for a direct transfer chemical shift assignments of natural abundance observable guanine (H1-N1, H21/H22-N2), thymine (H3-N3), and adenine (H61/H62-N6) cross peaks. Additional assignments were transferred by using 2D ^1^H,^1^H-TOCSY (*dipsi2gpph19*, 120 ms mixing time) and 2D ^1^H,^1^H-NOESY (*noesyesgpph*, 80 ms mixing time) spectra. As before, the same sample prepared in 100% D_2_O was used to collect a 2D ^1^H,^1^H-NOESY (*noesyesgpph*, 80 ms mixing time) spectra to help complete the additional sugar and base ^1^H chemical shift assignments. The assigned ^1^H and ^15^N chemical shifts for dopamine-bound RKEC1 at 278 K were deposited in the BMRB under accession code 52689, and the exchangeable ^1^H chemical shifts are also listed in **Supplementary Table 3**.

For the purpose of structure determination, it was also necessary to determine the chemical shift assignments at 298 K. Most of the ^1^H chemical shifts assigned at 278 K were transferred to 298 K by using two 2D ^1^H,^1^H-NOESY spectra (*noesyesgpph*, 80 ms mixing time) collected on the same samples of 2 mM RKEC1 with 2.25 mM dopamine in 90% H_2_O/10% D_2_O and in 100% D_2_O. In addition, a 2D ^1^H,^1^H-TOCSY (*dipsi2gpph19*, 120 ms mixing time) was collected on the 100% D_2_O sample. To remove any ambiguity for some of the guanosine assignments, we supplemented our data with 90% H_2_O/10% D_2_O samples of 1.5 mM RKEC1 and 1.75 mM dopamine in which G21, G25, G28, G29, G31 or G33 were separately enriched with ^13^C and ^15^N. As before, we collected sensitivity-enhanced ^1^H,^15^N-HSQC with a ^15^N spectral width of 2500 Hz, centered on 150 ppm or 75 ppm, for detection of guanine base H1-N1 imine or H21/H22-N2 amine cross peaks, respectively, and ^1^H and ^13^C sugar resonances based on 2D ^1^H,^13^C-HSQC spectra (*hsqcetgpsisp2.2*, with a ^13^C spectral width of 30 ppm centered on 75 ppm). Due to the excellent quality of the NMR data in 100% D_2_O at 298 K, we were able to assign to near completion all of the sugar ^1^H and ^13^C chemical shifts by using a long-acquisition natural abundance ^1^H,^13^C-HSQC spectrum in D_2_O (*hsqcetgpsisp2.2*, with a ^13^C spectral width of 30 ppm centered on 75 ppm), as illustrated in **Supplementary Fig. S3b**. In addition to the above 2D ^1^H,^1^H-TOCSY, a 2D DQF-COSY (*cosydfgpph19*) spectrum was used to unambiguously assign H2’ and H2’’ where possible. The assigned ^1^H, ^13^C and ^15^N chemical shifts for dopamine-bound RKEC1 at 298 K were deposited in the BMRB under accession code 52690, and the non-exchangeable ^1^H chemical shifts are also listed in **Supplementary Table 3**.

### Initial structure determination using ARIA/CNS

A first ensemble of structures was calculated for dopamine-bound RKEC1 by using ARIA2.3/CNS1.2.(Brunger, 2007; Rieping et al., 2007) In order to set up the system, we first generated CNS topology and parameter files for dopamine using PRODRG.(Schüttelkopf & van Aalten, 2004; van Aalten et al., 1996) The atom names generated during this step were maintained in order to prevent ambiguity with existing CNS topology and parameter information. Distance restraints involving exchangeable ^1^H were obtained at 278 K from a 2D ^1^H-^1^H NOESY spectrum (Bruker pulse program *noesyesgpph*, 80 ms mixing time) using a sample of 2 mM RKEC1 with 2.25 mM dopamine, prepared in 20 mM sodium phosphate (pH 7.4), 140 mM NaCl, 2 mM MgCl_2_ with 90% H_2_O/10% D_2_O. The same sample in 100% D_2_O was used to collect another 2D ^1^H,^1^H-NOESY spectrum (*noesyesgpph*, 80 ms mixing time) at 298 K, to obtain distance restraints involving non-exchangeable ^1^H. The initial 278 K peak list had all but 7 cross peaks out of 400 assigned, whereas the initial 298 K peak list had complete assignments for 818 out of 1263 cross peaks. From the chemical shift assignments, all nucleotides had an *anti* conformation except for G31 with downfield C8 and H1′ chemical shifts, an upfield H8 chemical shift, and a strong H8-H1′ NOE cross peak, consistent with a *syn* conformation. These dihedral restraints were included in the structure calculation. Based on the intensities of the H2′-H1′ and H2″-H1′ ^1^H,^1^H-TOCSY and DQF-COSY cross peaks during the chemical shift assignment, we also included dihedral restraints for 3′-endo sugar puckers for G17, T18, C20 and G34 instead of the default 2′-endo parameters for DNA. The final ensemble refined in explicit water consists of the 15 lowest energy structures from a total of 100 calculated models. Complete refinement statistics for this step are presented in **Table 1**.

### AMBER system preparation

In order to generate the final ensembles, we used the 15 models from ARIA/CNS as starting structures for restrained molecular dynamics in AMBER. The system was prepared using the LEAP program from the Amber24 suite.(Case et al., 2023) The OL21 and AMBER General Force Field for organic molecules (Version 1.81) were used to describe the complex.(Love et al., 2023; Wang et al., 2004; Zgarbová et al., 2021) The structure was explicitly solvated in a truncated octahedral box of water molecules, using the OPC model with the ad hoc Li/Merz ion parameters of atomic ions (12-6 set),(Izadi et al., 2014; Li et al., 2020) with a minimum of 14 Å between the solute and the box edge. For simulations in absence of salts (case of unrestrained simulations), the system was neutralized by adding 25 Na^+^ counter-cations. In cases where 140 mM NaCl were added to better reflect the SELEX conditions (restrained simulations), the number of required ions (𝑁_±_; here around 35 Na^+^ and 10 Cl^-^) was determined following the SLTCAP method,(Schmit et al., 2018) using Equation 1 simplified by Machado and Pantano (Machado & Pantano, 2020) into Equation 2, where ν_𝑤_ is the water volume of the simulation box (around 3 x 10^5^ Å^3^ here) in reduced units, 𝑐_0_ the salt concentration, 𝑄 the total charge of the complex (here-25: 26 phosphates on the aptamer and 1 ammonium group on the dopamine), and 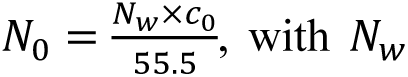 the number of water molecules in the simulation box (here, around 9 x 10^4^)(Machado & Pantano, 2020)

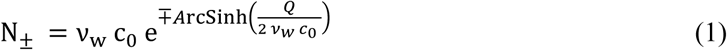

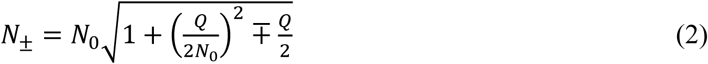

Note that the SPLIT method described by Machado and Pantano cannot be applied as our system does not satisfy the 𝑁_0_≫ 𝑄 condition; however it yields identical values in most cases or deviates by a single ion.

### Restrained minimization, heating and molecular dynamics

Using an in-house script based on R (https://www.R-project.org), the unambiguous distance restraints obtained in ARIA/CNS were converted to an 8-column format suitable for its processing by the makeDIST_RST function from the sander module of AMBER. A custom map file was prepared to define common names for groups of protons sharing given restraints (e.g. 3 hydrogens from the same methyl group). The resulting DISANG restraints file was then applied to all steps below.

All simulation steps were performed with pmemd.cuda (v. 18.0.0) from the CUDA version of AMBER,(Götz et al., 2012; Le Grand et al., 2013; Salomon-Ferrer et al., 2013) on an NVIDIA H100 PCIe Tensor core GPU (CUDA version: 12.4) from the DOREMI CALI v3 cluster of the *Mésocentre de Calcul Intensif Aquitain* (Univ. Bordeaux). The system was minimized for 20000 cycles using the steepest descent algorithm for the first 4000 steps and the conjugate gradient for the next 16000 steps. The weights of the restraints were kept constant at 100 kcal·mol^-1^·Å^-2^. The system was then heated at constant volume from 0 to 298 K over 18 ps then kept for 2 ps at the final temperature, using a time step of 2 fs, the Langevin thermostat with a 2.0 ps^-1^ collision frequency and a different seed for the pseudo-random number generation for every run to avoid synchronization artifacts,(Sindhikara et al., 2009) an 8 Å non-bonded cutoff, and the bonds involving hydrogen were constrained with the SHAKE algorithm. The system was further equilibrated five times at 298 K with the parameters above and the restraint weights ramping down from 100 to 5 kcal·mol^-1^·Å^-2^. The pressure was kept at 1 bar with the Berendsen barostat.(Berendsen et al., 1984) Restrained molecular dynamics for subsequent minimization were run for 10 ns with the parameters above, and restraints weights set at 20 kcal·mol^-1^·Å^-2^. The final coordinates were further minimized, as described above except for the restraint weights set at 20 kcal·mol^-1^.Å^-2^. Remaining NMR violations were summarized with the sviol function from the Amber package. Complete AMBER refinement statistics for the final ensemble of 15 structures is presented in **Table 1**. The ensemble has been deposited in the PDB as accession 9HIO.

Production MD were run over a microsecond with and without restraints, as well as in absence of dopamine, with the parameters above. Four replicates were generated for the restrained production run: two from the lowest-energy conformer identified by ARIA/CNS, and two from the most structurally distinct conformers selected by PCA (**Supplementary Fig. 6b**). Simulation stability was evaluated by computing the pairwise RMSD matrix across trajectory frames (**Supplementary Fig. 8**). Dihedral angles (χ, **Supplementary Figs. 9, 12**; sugar puckers, **Supplementary Figs. 10, 13**) and nucleotide H-bonding (**Supplementary Figs. 11, 14**) were tracked as described below.

## Structure analysis

The bio3d package was used for minimized structure and trajectory files cleanup, alignment, filtering, averaging and analysis.(Grant et al., 2006) The determination of RMSD and RMSF was performed with the rmsd and rmsf functions. Dihedral angles, sugar pucker angles and amplitudes 𝜃_𝑀_ were obtained with in-house R functions leveraging the torsion.xyz function, following Equation 3, where the pucker 𝑃 is determined by Equation 4 and the sugar torsion angles v_i_are defined by four atoms as shown below.

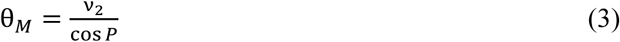

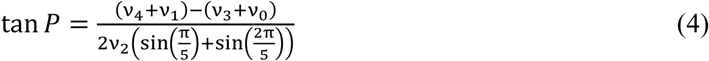

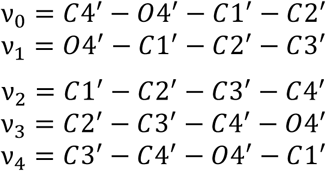

All atom-atom and ring-ring distances and angles were measured with the dist.xyz and angle.xyz functions. In-house R scripts were used to infer the formation of H-bonds from these values on full trajectories, by first selecting realistic donor/acceptor atoms for each residue/ligand, then verifying that the donor/acceptor distances and angles were compatible with H-bond formation.

Base pairs were further characterized with DSSR 2.3.2,(Lu et al., 2015) (Lu, 2020) using the web API through the httr2 R package (https://CRAN.R-project.org/package=httr2).

Principal component analysis on the final structures and trajectories (all atoms except hydrogen) was performed with the pca function from bio3d. The results were clustered with the k-means method using Euclidean distance and the best number of clusters determined by Nbclust.(Charrad et al., 2014) Cross-correlation analysis was carried out with the dccm function of bio3d.

Molecular structures images were created in PyMOL 2.5 (Schrödinger, LLC 2021, http://www.pymol.org/pymol), and ligand binding site diagram with LigPlot^+^.(Laskowski & Swindells, 2011) All further data processing and all plotting was performed in R 4.3. Apache Arrow (https://github.com/apache/arrow/) was used to write/read processed data files in feather format.

### DNA for fabrication of E-AB sensors

Aptamer sequences employed for the fabrication of E-AB sensors were purchased from Bio Basic Inc with HPLC purification. Sequences and modifications are shown in **Supplementary Table 6**. We used the oligonucleotides without further purification with resuspension at 100 µM in deionized water, determined using the molar absorption coefficient at 260 nm provided by the supplier with an Implen NP80 nanophotometer, prior to aliquoting 2 µL into separate tubes. BioBasic Inc chemically modified the 5′ end of aptamers with a thiol on a 6-carbon linker and the 3′ end with a carboxy-modified methylene blue attached to DNA via the formation of an amide bond to a primary amine on a 6-carbon linker.

### Electrode preparation

Prior to sensor fabrication, we mechanically cleaned the electrodes (2.0 mm diameter, CH Instruments) first by polishing them with microcloth pads soaked with a 1 μm diamond suspension oil slurry (Buehler), followed by sonication of the electrodes in ethanol for 5 min. A second mechanical cleaning step used 0.05 μm alumina oxide powder aqueous solution (Buehler). followed by sonication in distilled water for 5 min. We then cleaned the electrodes electrochemically through successive cathodic and anodic scans in NaOH and H_2_SO_4_ solutions. For electrochemical cleaning we relied on a CH Instruments 1040C multichannel potentiostat, using a standard three-electrode cell containing a platinum counter electrode with an Ag/AgCl reference electrode (CH Instruments). First, the electrodes were cleaned in 0.5 M NaOH by performing repeated cyclic voltammetry scans in the potential window from-1 to-1.6 V (all potentials versus Ag|AgCl) at the scan rate of 1 V s^-1^ for 300 cycles. Next, we moved the electrodes into a 0.5 M H_2_SO_4_ solution and performed a 2 V oxidizing chronoamperometric step for 5 s. Subsequently, we applied a reducing potential of-0.35 V for 10 s followed by voltammetric cycles at a scan rate of 4 V s^-1^ from-0.35 to 1.5 V for 10 scans and then two cycles using the same potential range at 0.1 V s^-1^. Finally, we washed the electrodes in deionized water and determined their surface area by acquiring a cyclic voltammogram from-0.35 to 1.5 V at 0.1 V s^-1^ in a 0.05 M H_2_SO_4_ solution and integrating the gold reduction peak.(Dauphin-Ducharme et al., 2022; Xiao et al., 2007)

### Sensor preparation

We first reduced the aptamer by incubating a 2 µL aliquot for approximately 1 h with a 4 µL volume of 10 mM aqueous tris(2-carboxyethyl)phosphine (TCEP, Sigma Aldrich) at room temperature in the dark. Following this step, we diluted the reduced aptamers in 1X PBS (Sigma Aldrich) to obtain a final concentration of 200 nM. Next we immersed the electrochemically cleaned gold electrodes in the 200 nM aptamer solution for 1 h in the dark. We rinsed the electrodes with 1X PBS and incubated for 3 h at room temperature in 1X PBS containing 5 mM 6-mercaptohexanol (Sigma Aldrich), followed by a rinse using 1X PBS prior to their use.

### Sensor characterization

We carried out electrochemical impedance spectroscopy measurements using a BioLogic VMP-300 potentiostat. For all experiments, we first performed a cyclic voltammogram from 0 to −0.5 V *vs* Ag/AgCl to determine the methylene blue redox potential via the average of the reduction and oxidation peak potentials in the binding buffer (1X PBS, 2 mM MgCl_2_ containing 0.02% (w/v) ascorbic acid). Following this, we utilized this potential for potentiostatic electrochemical impedance measurements as a fixed value and applied a sinusoidal perturbation of 10 mV in amplitude. We then varied the frequencies of the oscillations from 10 kHz to 0.1 Hz and carried out 10 measurements per decade of frequencies. We repeated these measurements by exposing the sensors to increasing concentrations of dopamine in the binding buffer. We performed a nonlinear fit of the obtained impedance traces using BioLogic’s algorithms. From this, we determined the charge-transfer resistance (i.e., 𝑅_𝑐𝑡_) related to the electron-transfer reaction of the redox reporter. We then plotted the difference in measured charge-transfer resistance with the one measured in absence of dopamine against its concentrations to a Langmuir-Hill equation (Equation 5).(Nguyen et al., 2024; Rahbarimehr et al., 2023)

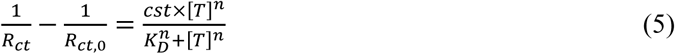

Where normalized 𝑅_𝑐𝑡,0_ and 𝑅_𝑐𝑡_ represents the minimal and measured charge transfer resistance, [𝑇] is the dopamine concentration, 𝑐𝑠𝑡 is a parameter combining electron transfer kinetics of the redox reporter and total concentration of aptamer on the surface, 𝑛 is the Hill coefficient, 𝐾_𝐷_ is the dissociation constant.

## Data availability

All data supporting the findings of this work are available within the main text and its Supplementary Information. The list of oligonucleotides used in this work is provided in **Supplementary Table 2**. Chemical shift assignments are available from the BMRB under accession number 52688 for DA-mut3-del7, and accession numbers 52689 and 52690 for dopamine-bound RKEC1 at 278 K and 298 K, respectively. The ensemble of 15 structures of dopamine-bound RKEC1 were deposited in the PDB under accession number 9HIO.

## Acknowledgments

We thank Estelle Morvan, Axelle Grélard, and the structural biophysical chemistry facility at the European Institute of Chemistry and Biology (CNRS UAR 3033, Inserm US001) for access to NMR spectrometers.

## Funding and additional information

Financial support from the IR INFRANALYTICS FR2054 for conducting the research is gratefully acknowledged. This work was supported by funding from the Fonds de Recherche du Québec, Nature et Technologies, and the Natural Sciences and Engineering Research Council of Canada (NSERC) to P.D.-D. and P.E.J. through the NOVA program, and a France-Canada Research Grant to C.D.M. and P.E.J.

## Contributions

**C.D.M.**, **P.E.J.** and **P.D.-D.** conceptualized the project and acquired funding. **H.P.C.**, **Y.A.K.** and **P.E.J.** performed and analyzed the ITC experiments. **B.V.** synthesized and purified the isotope-enriched DNA. **C.D.M.** prepared the NMR samples, collected spectra, and analyzed the data. **C.D.M.** and **E.L.** determined the structure. **E.L.** performed the molecular dynamics simulations, performed computation analysis, and is responsible for data curation. **M.-D.N.** and **P.D.-D.** designed, performed and analyzed the E-AB biosensor studies. **C.D.M.**, **P.E.J.** and **P.D.-D.** wrote the original draft with review and editing by all authors. All authors approved the final version.

