## Supplementary Material for "Unconventional DNA architecture in a dopamine–bound aptamer complex"

**Supplementary Table 1 | ITC data for DA-mut3 and RKEC1 binding to dopamine and related ligands.**[a]

| Aptamer | Ligand | $K_d$ (μM) | ΔH (kcal/mol) | -TΔS (kcal/mol) |
|---|---|---|---|---|
| DA-mut3 | Dopamine | 0.8 ± 0.1 | -27 ± 1 | 19 ± 1 |
| | Tyramine | 76 ± 7 | -13 ± 0 | 8 ± 0.1 |
| | 6-Hydroxydopamine [b] | 0.7 ± 0.1 | -32 ± 4 | 24 ± 4 |
| | Dopamine [b] | 0.9 ± 0.1 | -34 ± 3 | 26 ± 3 |
| | 3-hydroxy-4-methoxyphenethylamine | 118 ± 5 | -12 ± 0.7 | 7 ± 0.7 |
| | 3-Methoxytyramine | 700 ± 100 | -4 ± 0.7 | -0.2 ± 0.5 |
| | Norepinephrine | 6 ± 0.2 | -25 ± 0.2 | 18 ± 0.2 |
| | Epinephrine | 29 ± 4 | -26 ± 7 | 19 ± 7 |
| | 3,4-dihydroxybenzylamine | 6 ± 0.1 | -26 ± 4 | 19 ± 4 |
| | Catechol /1,2 dihydroxy benzene | *NB* | | |
| | 3,4-dihydroxyhydrocinnamic acid | *NB* | | |
| | 3,4 dihydroxyphenylacetic acid (DOPAC) | *NB* | | |
| | Tyrosine | *NB* | | |
| | Phenylalanine | *NB* | | |
| DA-mut3-del7 | Dopamine | 108 ± 3 | -41 ± 1 | 36 ± 1 |
| RKEC1 | Dopamine | 4.0 ± 0.6 | -43 ± 7 | 36 ± 7 |
| | Tyramine | 800 ± 100 | -32 ± 4 | 28 ± 4 |
| | 3,4-dihydroxybenzylamine | 72 ± 2 | -25 ± 0.6 | 19 ± 0.7 |
| | Norepinephrine | 46 ± 1 | -24 ± 0.2 | 18 ± 0.2 |

[a] Data acquired at 20 °C in 20 mM $Na_xH_yPO_4$, 140 mM NaCl, 2 mM $MgCl_2$, pH 7.4. The values reported are averages of two individual experiments. *NB* denotes no binding.

[b] Measured in the same buffer plus 1% ascorbic acid as required for 6-hydroxydopamine stability

**Supplementary Table 2 | Oligonucleotide sequences**

| Name | Sequence | Notes |
|---|---|---|
| DA-mut3 | CGACGTCAGTTTGAAGGTTCGTTCGCAGGTGTGGAGTGACGTCG | ITC, NMR, reference for binding |
| DA-mut3-del7 | AGTTTGAAGGTTCGTTCGCAGGTGTGGAGT | ITC, NMR, initial aptamer for structure determination |
| DA-mut3-del7-*G9 | AGTTTGAAGGTTCGTTCGCAGGTGTGGAGT | * = 5%-$^{13}C^{15}N$, NMR assignment |
| DA-mut3-del7-*G13 | AGTTTGAAGGTTCGTTCGCAGGTGTGGAGT | * = 5%-$^{13}C^{15}N$, NMR assignment |
| DA-mut3-del7-*G16 | AGTTTGAAGGTTCGTTCGCAGGTGTGGAGT | * = 5%-$^{13}C^{15}N$, NMR assignment |
| DA-mut3-del7-*G17 | AGTTTGAAGGTTCGTTCGCAGGTGTGGAGT | * = 5%-$^{13}C^{15}N$, NMR assignment |
| DA-mut3-del7-*G21 | AGTTTGAAGGTTCGTTCGCAGGTGTGGAGT | * = 5%-$^{13}C^{15}N$, NMR assignment |
| DA-mut3-del7-*G25 | AGTTTGAAGGTTCGTTCGCAGGTGTGGAGT | * = 5%-$^{13}C^{15}N$, NMR assignment |
| DA-mut3-del7-*G28 | AGTTTGAAGGTTCGTTCGCAGGTGTGGAGT | * = 5%-$^{13}C^{15}N$, NMR assignment |
| DA-mut3-del7-*G29 | AGTTTGAAGGTTCGTTCGCAGGTGTGGAGT | * = 5%-$^{13}C^{15}N$, NMR assignment |
| DA-mut3-del7-*G31 | AGTTTGAAGGTTCGTTCGCAGGTGTGGAGT | * = 5%-$^{13}C^{15}N$, NMR assignment |
| DA-mut3-del7-*G33 | AGTTTGAAGGTTCGTTCGCAGGTGTGGAGT | * = 5%-$^{13}C^{15}N$, NMR assignment |
| DA-mut3-del7-*G34 | AGTTTGAAGGTTCGTTCGCAGGTGTGGAGT | * = 5%-$^{13}C^{15}N$, NMR assignment |
| DA-mut3-del7-*G36 | AGTTTGAAGGTTCGTTCGCAGGTGTGGAGT | * = 5%-$^{13}C^{15}N$, NMR assignment |
| DA-mut3-del7-*T10 | AGTTTGAAGGTTCGTTCGCAGGTGTGGAGT | * = 5%-$^{13}C^{15}N$, NMR assignment, only 50% of desired oligo (NMR spectrum leads to RKEC1) |
| DA-mut3-del7-*T11 | AGTTTGAAGGTTCGTTCGCAGGTGTGGAGT | * = 5%-$^{13}C^{15}N$, NMR assignment |
| DA-mut3-del7-*T12 | AGTTTGAAGGTTCGTTCGCAGGTGTGGAGT | * = 5%-$^{13}C^{15}N$, NMR assignment |
| DA-mut3-del7-*T18 | AGTTTGAAGGTTCGTTCGCAGGTGTGGAGT | * = 5%-$^{13}C^{15}N$, NMR assignment |
| DA-mut3-del7-*T19 | AGTTTGAAGGTTCGTTCGCAGGTGTGGAGT | * = 5%-$^{13}C^{15}N$, NMR assignment |
| DA-mut3-del7-*T22 | AGTTTGAAGGTTCGTTCGCAGGTGTGGAGT | * = 5%-$^{13}C^{15}N$, NMR assignment |
| DA-mut3-del7-*T23 | AGTTTGAAGGTTCGTTCGCAGGTGTGGAGT | * = 5%-$^{13}C^{15}N$, NMR assignment |
| DA-mut3-del7-*T30 | AGTTTGAAGGTTCGTTCGCAGGTGTGGAGT | * = 5%-$^{13}C^{15}N$, NMR assignment |
| DA-mut3-del7-*T32 | AGTTTGAAGGTTCGTTCGCAGGTGTGGAGT | * = 5%-$^{13}C^{15}N$, NMR assignment |
| DA-mut3-del7-*A8 | AGTTTGAAGGTTCGTTCGCAGGTGTGGAGT | * = 5%-$^{13}C^{15}N$, NMR assignment |
| DA-mut3-del7-*A14 | AGTTTGAAGGTTCGTTCGCAGGTGTGGAGT | * = 5%-$^{13}C^{15}N$, NMR assignment |
| DA-mut3-del7-*A15 | AGTTTGAAGGTTCGTTCGCAGGTGTGGAGT | * = 5%-$^{13}C^{15}N$, NMR assignment |
| DA-mut3-del7-*A27 | AGTTTGAAGGTTCGTTCGCAGGTGTGGAGT | * = 5%-$^{13}C^{15}N$, NMR assignment |
| DA-mut3-del7-*A35 | AGTTTGAAGGTTCGTTCGCAGGTGTGGAGT | * = 5%-$^{13}C^{15}N$, NMR assignment |
| RKEC1 | TTGAAGGTTCGTTCGCAGGTGTGGAGT | ITC, NMR, aptamer used for full structure determination |
| RKEC1-*G21 | TTGAAGGTTCGTTCGCAGGTGTGGAGT | * = 5%-$^{13}C^{15}N$, NMR assignment |
| RKEC1-*G25 | TTGAAGGTTCGTTCGCAGGTGTGGAGT | * = 5%-$^{13}C^{15}N$, NMR assignment |
| RKEC1-*G28 | TTGAAGGTTCGTTCGCAGGTGTGGAGT | * = 5%-$^{13}C^{15}N$, NMR assignment |
| RKEC1-*G29 | TTGAAGGTTCGTTCGCAGGTGTGGAGT | * = 5%-$^{13}C^{15}N$, NMR assignment |
| RKEC1-*G31 | TTGAAGGTTCGTTCGCAGGTGTGGAGT | * = 5%-$^{13}C^{15}N$, NMR assignment |
| RKEC1-*G33 | TTGAAGGTTCGTTCGCAGGTGTGGAGT | * = 5%-$^{13}C^{15}N$, NMR assignment |

**Supplementary Table 3 | $^{1}$H Chemical shift assignments for dopamine-bound RKEC1.**[a]

| | 278 K | | 298 K | | | | | |
|---|---|---|---|---|---|---|---|---|
| Residue | Gua-H1 Thy-H3 | Gua-H21/H22 Cyt-H41/H42 Ade-H61/H62 | H1′ | H2′ / H2″ | H3′ | H4′ | H5′ / H5″ | Base |
| T11 | 10.985 | | 6.358 | 1.919 / 2.928 | 4.933 | 4.238 | 3.952 / 3.827 | 7.760 (H6), 1.609 (M7) |
| T12 | 14.181 | | 5.834 | 2.298 / 2.632 | 5.020 | 4.307 | 4.048 / - | 7.442 (H6), 1.757 (M7) |
| G13 | 10.302 | - / 6.833 | 5.880 | 2.802 / 2.698 | 5.067 | 4.500 | 4.320 / 4.250 | 8.027 (H8) |
| A14 | | - / - | 6.053 | 2.163 / 2.428 | 4.566 | - | 3.854 / 3.214 | 7.658 (H2), 8.059 (H8) |
| A15 | | 7.678 / 9.078 | 6.247 | 2.769 / 3.009 | 4.855 | 4.401 | 4.078 / 3.802 | 8.014 (H2), 8.281 (H8) |
| G16 | 13.130 | 7.645 / 6.519 | 4.914 | 2.561 / 2.651 | 4.992 | 4.398 | 4.220 / - | 7.986 (H8) |
| G17 | 14.218 | 6.525 / 7.513 | 6.215 | 2.694 / 2.694 | 4.872 | 4.393 | 4.355 / 4.296 | 8.052 (H8) |
| T18 | 9.582 | | 6.171 | 2.172 / 2.389 | 4.819 | 4.303 | 4.382 / 4.207 | 7.435 (H6), 1.173 (M7) |
| T19 | 12.541 | | 6.294 | 2.292 / 2.807 | 4.888 | 4.353 | 4.280 / 4.191 | 7.446 (H6), 1.504 (M7) |
| C20 | | 7.767 / 6.613 | 5.951 | 2.079 / 1.384 | 4.871 | 3.819 | 3.963 / 4.062 | 5.848 (H5), 7.397 (H6) |
| G21 | 13.781 | 9.107 / 5.035 | 6.037 | 2.989 / 2.596 | 5.046 | 4.480 | 4.106 / - | 8.098 (H8) |
| T22 | - | | 6.178 | 1.946 / 2.271 | 4.737 | 3.752 | 3.805 / 3.742 | 7.560 (H6), 1.641 (M7) |
| T23 | - | | 6.441 | 2.320 / 2.497 | 4.483 | 4.557 | 4.164 / 3.992 | 7.788 (H6), 1.891 (M7) |
| C24 | | 7.898 / 8.609 | 6.233 | 1.747 / 2.134 | 4.871 | 4.333 | 4.062 / 3.971 | 6.165 (H5), 7.619 (H6) |
| G25 | - | - / - | 7.032 | 3.169 / 2.873 | 5.269 | 4.862 | 4.325 / 4.060 | 8.234 (H8) |
| C26 | | 6.961 / 8.220 | 5.700 | -0.386 / 1.337 | 3.991 | 4.078 | 3.956 / 3.815 | 5.640 (H5), 7.684 (H6) |
| A27 | | 7.082 / 8.336 | 3.915 | 2.802 / 2.541 | 4.789 | 4.300 | 3.772 / - | 8.179 (H2), 8.301 (H8) |
| G28 | 11.606 | 6.248 / 9.716 | 6.171 | 2.504 / 2.756 | 5.005 | 4.557 | 4.140 / 4.066 | 7.285 (H8) |
| G29 | 13.815 | 7.870 / 7.518 | 5.672 | 2.752 / 2.391 | 5.016 | 4.428 | 4.200 / 4.579 | 8.256 (H8) |
| T30 | - | | 5.982 | 1.718 / 2.086 | 4.558 | 3.083 | 3.357 / 2.773 | 7.395 (H6), 1.525 (M7) |
| G31 | 11.340 | 5.969 / 6.762 | 5.992 | 3.497 / 2.667 | 4.714 | 4.304 | 3.941 / - | 7.630 (H8) |
| T32 | 9.953 | | 5.641 | 2.254 / 2.254 | 4.895 | 4.140 | 4.044 / 4.139 | 7.280 (H6), 1.369 (M7) |
| G33 | 12.631 | - / - | 5.924 | 2.804 / 3.051 | 5.009 | 4.200 | 4.014 / 3.943 | 7.524 (H8) |
| G34 | 11.631 | - / 5.799 | 5.524 | 2.554 / 2.320 | 4.915 | 4.277 | 4.202 / - | 7.673 (H8) |
| A35 | | 8.432 / 8.124 | 5.899 | 1.412 / 2.060 | 4.779 | 4.296 | 4.093 / 4.023 | 8.085 (H2), 7.382 (H8) |
| G36 | 7.701 | - / - | 5.473 | 2.791 / 2.480 | 5.011 | 4.313 | 3.877 / 3.877 | 7.990 (H8) |
| T37 | - | | 5.939 | 2.231 / 2.063 | 4.370 | 4.126 | 4.005 / 4.005 | 7.482 (H6), 1.644 (M7) |

[a] Additional $^{13}$C and $^{15}$N chemical shift assignments can be found in BMRB accession numbers 52689 (278 K) and 52690 (298 K).

**Supplementary Table 4 | Intermolecular dopamine–RKEC1 restraints.**

| Ligand | DNA | Restraint (Å) |
|---|---|---|
| Dopamine H2 | T18 H3 | 4.0 ± 2.0 |
| | T19 H3 | 3.6 ± 1.6 |
| | T19 M7 | 4.7 ± 2.7 |
| | A27 H2 | 1.6 ± 0.3 |
| | T32 H3 | 4.4 ± 2.4 |
| | T32 M7 | 4.0 ± 2.0 |
| | G33 H1 | 3.2 ± 1.3 |
| Dopamine H5 | T32 H1′ | 3.4 ± 1.4 |
| | T32 H2′ | 4.2 ± 2.2 |
| | T32 H2″ | 3.7 ± 1.7 |
| | T32 H3 | 4.7 ± 2.7 |
| | G33 H5′ | 4.7 ± 2.8 |
| | G33 H5″ | 2.8 ± 1.0 |
| | G33 H8 | 4.9 ± 3.0 |
| Dopamine H6 | T32 H3 | 4.5 ± 2.6 |
| | G33 H1 | 4.4 ± 2.4 |
| Dopamine H71 | T19 H3 | 5.3 ± 3.5 |
| | A27 H2 | 1.8 ± 0.4 |
| | T32 H3 | 5.2 ± 3.4 |
| | G33 H1 | 3.4 ± 1.4 |
| Dopamine H72 | A27 H2 | 3.8 ± 1.9 |
| | T32 H3 | 4.6 ± 2.7 |
| | G33 H1 | 3.8 ± 1.8 |
| Dopamine HB1 | T32 H3 | 2.7 ± 0.9 |
| | A27 H2 | 3.6 ± 1.6 |
| Dopamine HB2 | A27 H2 | 2.2 ± 0.6 |
| | G28 H1 | 5.1 ± 3.3 |
| | G28 H21 | 3.3 ± 1.4 |
| | T32 H3 | 2.5 ± 0.8 |
| | G33 H1 | 5.8 ± 4.2 |
| Dopamine MN1 | T32 H3 | 4.7 ± 2.8 |
| Dopamine HO1 | T18 H3 | 2.2 ± 0.6 |
| | T19 H3 | 4.0 ± 2.0 |
| | T19 M7 | 1.9 ± 0.4 |
| | C20 H41 | 4.2 ± 2.2 |
| | C26 H41 | 3.3 ± 1.4 |
| | C26 H42 | 3.4 ± 1.5 |
| | A27 H2 | 3.6 ± 1.6 |
| | T32 M7 | 2.9 ± 1.0 |
| | G33 H1 | 4.8 ± 2.0 |
| Dopamine HO2 | T18 H3 | 2.5 ± 0.8 |
| | T18 M7 | 3.5 ± 1.6 |
| | T32 M7 | 2.9 ± 1.1 |

**Supplementary Table 5 | Base pairs identified by DSSR in the ensemble of 15 structures.[1]**

| Nucleotide 1 | Nucleotide 2 | Base pair[a] | Saenger | Name | Leontis-Westhof | DSSR[b] | Occurrance[c] |
|---|---|---|---|---|---|---|---|
| T11 | G17 | T-G | | | cWS | cW-m | 15 |
| T11 | G34 | T+G | XXVII | rWobble | tWW | tW+W | 15 |
| T12 | A35 | T+A | XXI | rWC | tWW | tW+W | 15 |
| G13 | G36 | G+G | IV | | tSS | tm+m | 15 |
| A15 | A35 | A-A | V | | tWH | tW-M | 15 |
| G16 | G34 | G-G | VII | | tWH | tW-M | 15 |
| G17 | G33 | G-G | VII | | tWH | tW-M | 15 |
| G17 | G34 | G-G | VII | | tWH | tW-M | 15 |
| T19 | A27 | T+A | XXI | rWC | tWW | tW+W | 15 |
| C20 | G28 | C+G | XXII | rWC | tWW | tW+W | 15 |
| C20 | T32 | C-T | | | tW. | tW-. | 15 |
| G21 | G29 | G+G | IV | | tSS | tm+m | 15 |
| G21 | G31 | G+G | III | | tWW | tW+W | 15 |
| C24 | G29 | C-G | XIX | WC | cWW | cW-W | 15 |
| C26 | G33 | C-G | XIX | WC | cWW | cW-W | 15 |
| G28 | T32 | G-T | | | cSW | cm-W | 15 |
| T18 | G33 | T-G | | | tWH | tW-M | 12 |
| T12 | G16 | T-G | | | cSS | cm-m | 8 |
| G25 | G28 | G-G | | | tSH | tm-M | 8 |
| T22 | T30 | T+T | | | tHH | tM+M | 7 |
| A15 | G36 | A-G | | | cWW | cW-W | 3 |
| C24 | G28 | C-G | | | tWH | tW-M | 3 |
| G25 | G29 | G-G | | | cWW | cW-W | 3 |
| T18 | C26 | T+C | | | tW. | tW+. | 1 |

[a] abbreviations: +, paired nucleotides have the same faces; -, the paired nucleotides have opposite faces (as in canonical antiparallel Watson-Crick pairing); WC, Watson-Crick pairing; W: Watson-Crick edge; H, Hoogsteen edge; S, sugar edge; M, major groove edge; m, minor groove edge; r: reverse; c/t: cis/trans orientation along the glycosidic bond.

[b] a dot (.) indicates an undefined edge

[c] number of observations out of the ensemble of 15 structures

### Supplementary Table 6

| sequence name | 5' modification | sequence (5' to 3') | 3' modification | internal modification |
|---|---|---|---|---|
| RKEC1 – MB regular | HO-$(CH_2)_6$-S-S-$(CH_2)_6$-O- | TTG AAG GTT CGT TCG CAG GTG TGG AGT | -O-$(CH_2)_6$-NH-CO-$(CH_2)_2$-MB | - |
| RKEC1 – MB inverse | -O-$(CH_2)_6$-NH-CO-$(CH_2)_2$-MB | TTG AAG GTT CGT TCG CAG GTG TGG AGT | HO-$(CH_2)_6$-S-S-$(CH_2)_6$-O- | - |
| RKEC1 – MB internal | - | TTG **T**(MB) AGG TTC GTT CGC AGG TGT GGA GT | HO-$(CH_2)_6$-S-S-$(CH_2)_6$-O- | **Thymine**-O-$(CH_2)_6$-NH-CO-$(CH_2)_2$-MB |

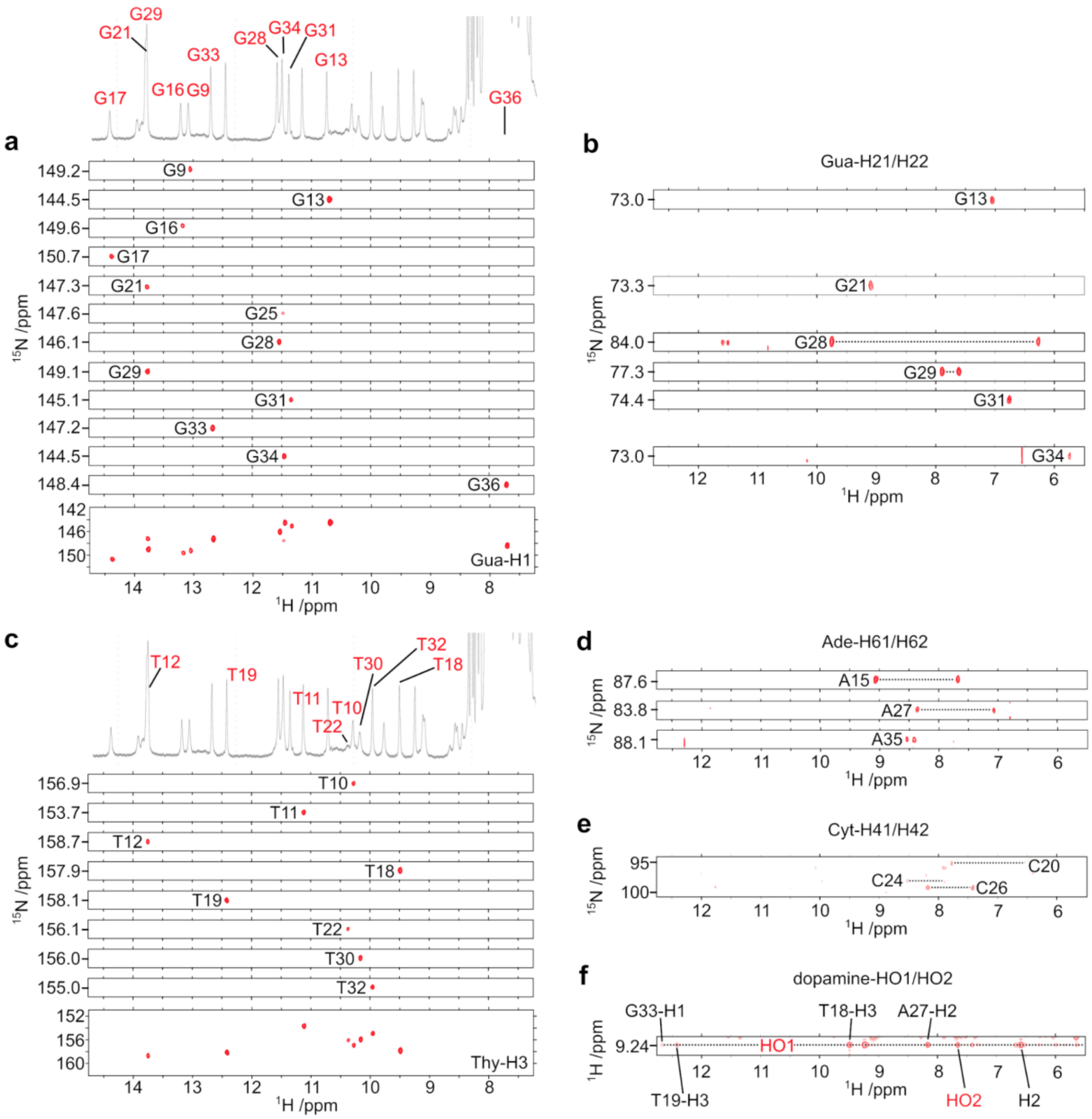

**Supplementary Fig. 1 | Unambiguous assignment of exchangeable $^1$H resonances for the dopamine-bound DA-mut3-del7**. Spectra were collected at 278 K at a field strength of 800 MHz, with samples at 2 mM in a buffer of 20 mM sodium phosphate (pH 7.4), 140 mM NaCl, 2 mM $MgCl_2$ and 10% $D_2O$. (**a**) N1-H1 imino cross peaks from $^1$H,$^{15}$N-HSQC spectra of the 12 guanosine-labeled DA-mut3-del7 complexes. (**b**) Observable H21/H22-N2 guanosine amino cross peaks from the same guanosine-labeled DA-mut3-del7 samples. (**c**) N3-H3 imino cross peaks from $^1$H,$^{15}$N-HSQC spectra are observed for 8 of the 10 thymidine-labeled DA-mut3-del7 complexes. There were no observed H3-N3 cross peaks for the T23 and T37 labeled samples. (**d**) Observable N6-H61/H62 adenosine amino cross peaks from the adenosine-labeled DA-mut3-del7 complexes. (**e**) All three cytidines have observable H41-N4 and H42-N4 amino cross peaks that were later assigned using an $^1$H,$^{15}$N-HSQC spectrum on a natural abundance sample, and comparison to the dopamine-bound RKEC1 data. (**f**) The bound dopamine hydroxyl $^1$H at 9.19 ppm (HO1) and 7.87 ppm (HO2) were later assigned based on a $^1$H-$^1$H NOESY spectrum and comparison to the dopamine-bound RKEC1 data.

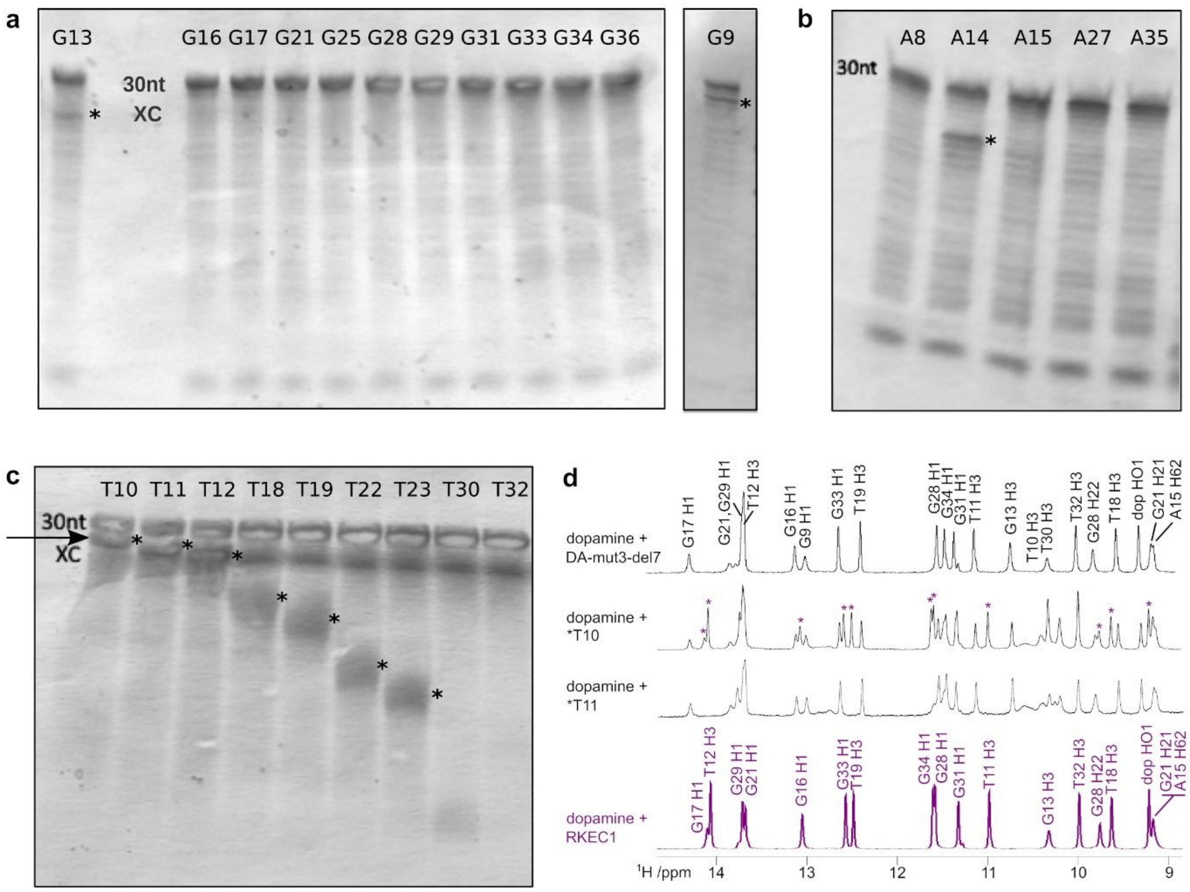


**Supplementary Fig. 2**. | **Chemical synthesis of residue-specific 5% $^{13}$C,$^{15}$N-labelled DA-mut3-del7**. (**a**,**b**) Site-specific incorporation of (a) $^{13}$C,$^{15}$N-guanosine phosphroamidite and (b) $^{13}$C,$^{15}$N-adenosine phosphoramidite produced samples with predominantly full-length oligonucleotides. (**c**) Significant amounts of 5′-abortive oligonucleotides were produced when using $^{13}$C,$^{15}$N-thymidine phosphoramidites, during the 3′ to 5′ chemical synthesis. These truncated oligonucleotides (indicated with an asterisk) stopped at the nucleotide 3' of the desired nucleotide targeted for isotope enrichment. Denaturing polyacrylamide gels were visualized with StainsAll, and the position of the loading dye xylene cyanol is indicated (XC). (**d)** In contrast to the second species present in the sample with abortive oligonucleotides at the T10 position, all other truncated oligonucleotides were unable to form an observable complex, as illustrated by the abortive oligonucleotides at position T11. The NMR spectra for dopamine-bound DA-mut3-del7, DA-mut3-del7-T10* and RKEC1 are from **Fig. 1d-f**.

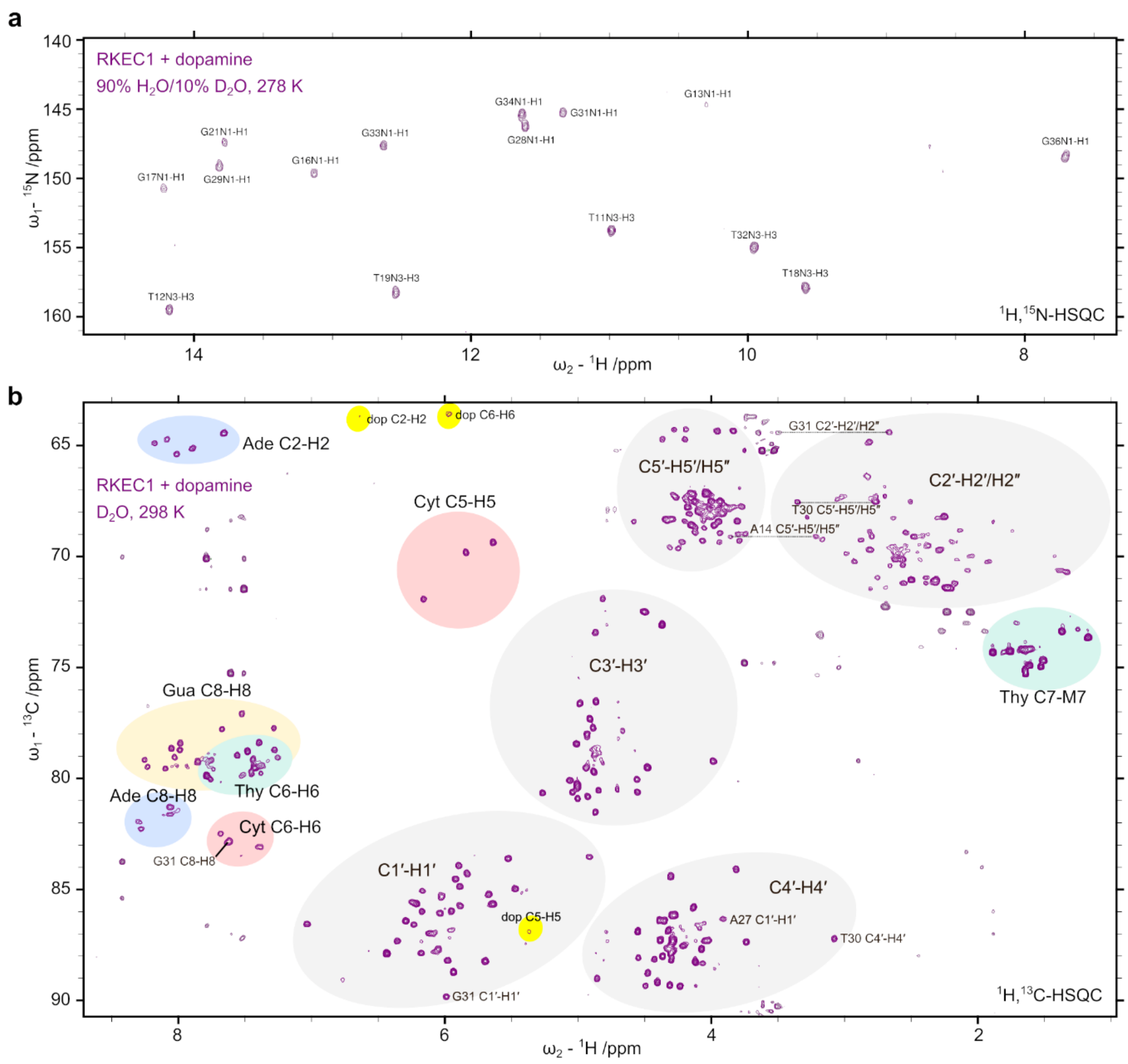


**Supplementary Fig. 3 | Annotated spectra for dopamine-bound RKEC1.** (**a**) Natural abundance $^{1}H,^{15}N$-HSQC spectrum at 278 K for a sample of 2 mM RKEC1 with 2.25 mM dopamine in a buffer of 20 mM sodium phosphate (pH 7.4), 140 mM NaCl, 2 mM $MgCl_2$ and 10% $D_2O$, acquired at a field strength of 800 MHz. Guanine N1-H1 and thymine N3-H3 cross peaks are annotated. (**b**) Natural abundance $^{1}H,^{13}C$-HSQC spectrum of 2 mM RKEC1 with 2.25 mM in the same buffer as in (a) but with 100 % $D_2O$, using acquisition at 298 K and a field strength of 800 MHz. The spectrum allows for aliasing in the $\omega_1$ dimension to capture all $^{13}C$-$^{1}H$ cross peaks in a single spectrum. Atom types are grouped and annotated by colored circles, with outlier cross peaks individually annotated. Also note the downfield C8 and H1′ chemical shifts, and upfield H8 chemical shift, of G31, consistent with a *syn* conformation.

**a**

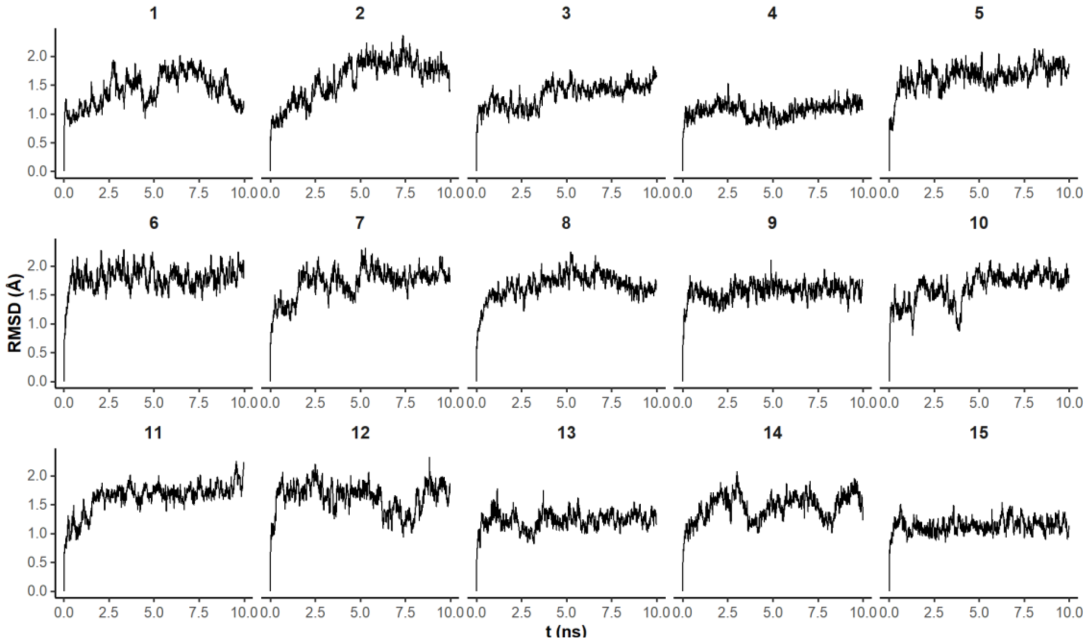


**b**

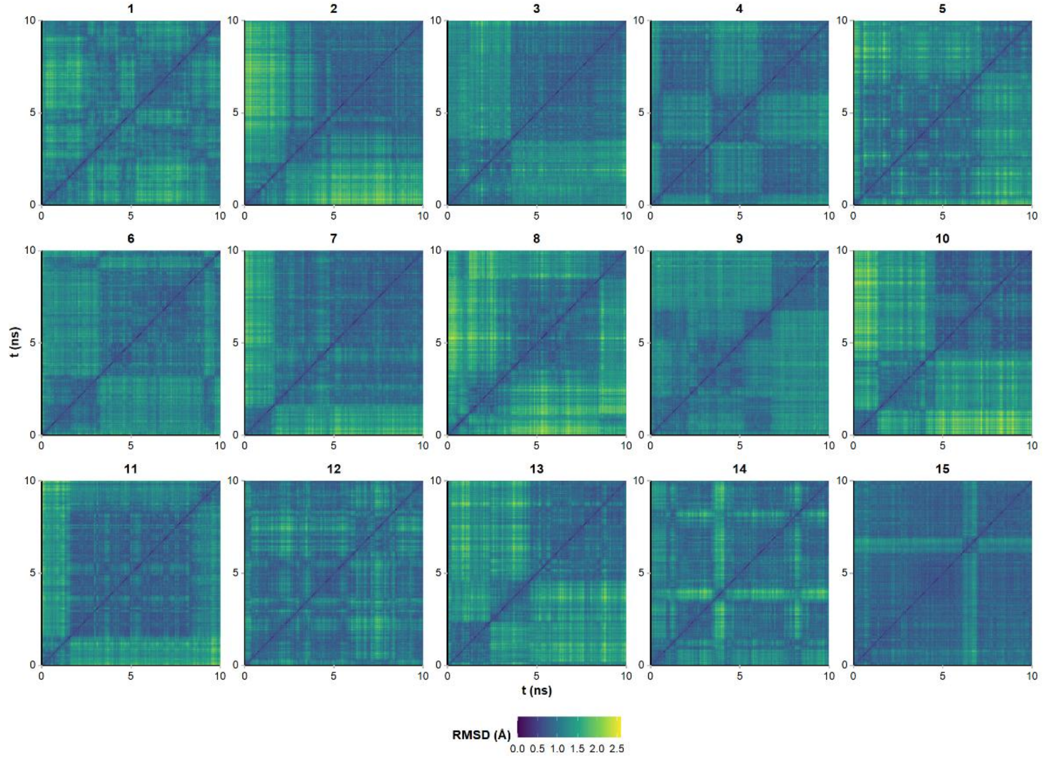


**Supplementary Fig. 4 | RMSD analysis during the 15 restrained MD simulations to generate to ensemble.** (a) Heavy-atom RMSD of all residues over the 10 ns restrained trajectories, computed with the first frame as reference. Each panel corresponds to one model. The first frame is used as reference. (**b**) Full pairwise heavy-atom RMSD matrices for the trajectories shown as heatmaps. Square regions along the diagonal indicate intervals of structural similarity within each trajectory.

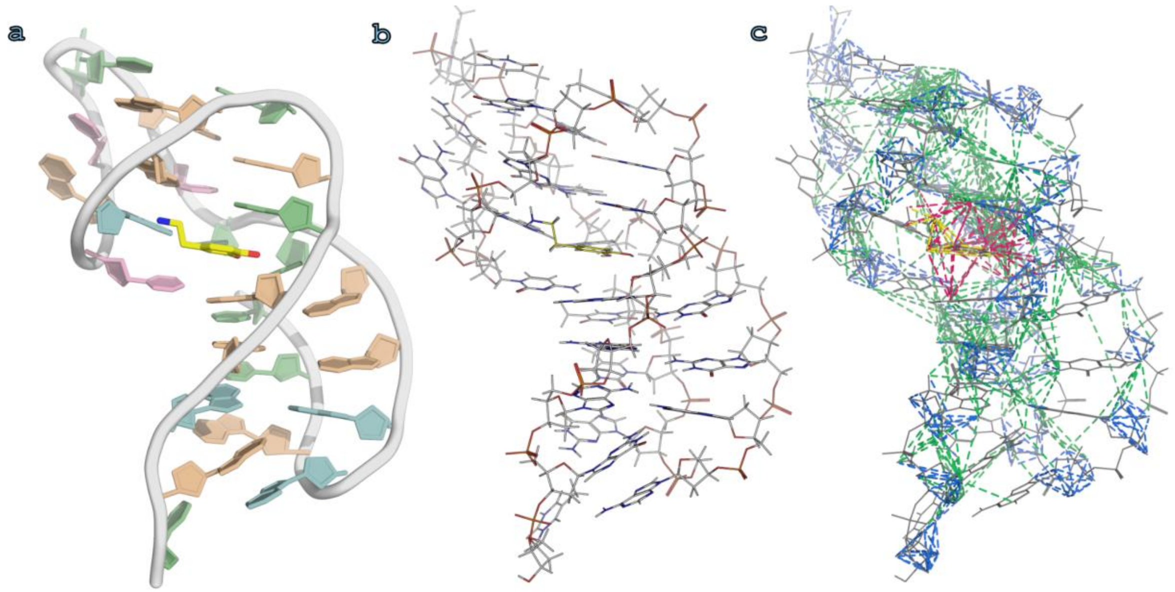


**Supplementary Fig. 5 | Distance restraints visualized on a representative model from the ensemble**. (a-**c**) Cartoon (a) and line (b) representation of the closest to average dopamine–RKEC1 structural model to aid in the interpretation of (c) which illustrates all distance restraints onto the line representation. Restrains are shown as dotted lines, colored blue for DNA intra-residue, green for DNA inter-residue, red for DNA–dopamine, and yellow for dopamine intermolecular restraints.

**a**

| State | 1 | 2 | 3 | 4 | 5 | 6 | 7 | 8 | 9 | 10 | 11 | 12 | 13 | 14 | 15 |
|---|---|---|---|---|---|---|---|---|---|---|---|---|---|---|---|
| 1 | | 1.30 | 1.56 | 1.48 | 1.61 | 1.80 | 1.69 | 1.40 | 1.59 | 1.63 | 1.83 | 1.38 | 1.59 | 1.28 | 1.58 |
| 2 | 1.30 | | 1.42 | 1.16 | 1.24 | 1.47 | 1.22 | 1.51 | 1.29 | 1.17 | 1.48 | 1.45 | 1.21 | 1.09 | 1.22 |
| 3 | 1.56 | 1.42 | | 1.75 | 1.11 | 1.18 | 0.98 | 1.42 | 1.25 | 1.07 | 1.11 | 1.40 | 1.01 | 1.34 | 1.20 |
| 4 | 1.48 | 1.16 | 1.75 | | 1.66 | 1.73 | 1.57 | 1.92 | 1.77 | 1.63 | 1.94 | 1.82 | 1.66 | 1.31 | 1.38 |
| 5 | 1.61 | 1.24 | 1.11 | 1.66 | | 0.96 | 0.94 | 1.38 | 1.22 | 0.97 | 0.91 | 1.19 | 0.90 | 1.24 | 1.10 |
| 6 | 1.80 | 1.47 | 1.18 | 1.73 | 0.96 | | 0.95 | 1.42 | 1.43 | 0.94 | 1.01 | 1.19 | 1.03 | 1.51 | 1.16 |
| 7 | 1.69 | 1.22 | 0.98 | 1.57 | 0.94 | 0.95 | | 1.56 | 1.30 | 0.72 | 1.06 | 1.43 | 1.00 | 1.15 | 1.11 |
| 8 | 1.40 | 1.51 | 1.42 | 1.92 | 1.38 | 1.42 | 1.56 | | 1.19 | 1.49 | 1.40 | 1.03 | 1.20 | 1.60 | 1.45 |
| 9 | 1.59 | 1.29 | 1.25 | 1.77 | 1.22 | 1.43 | 1.30 | 1.19 | | 1.25 | 1.30 | 1.41 | 1.04 | 1.36 | 1.28 |
| 10 | 1.63 | 1.17 | 1.07 | 1.63 | 0.97 | 0.94 | 0.72 | 1.49 | 1.25 | | 1.01 | 1.31 | 0.99 | 1.26 | 1.17 |
| 11 | 1.83 | 1.48 | 1.11 | 1.94 | 0.91 | 1.01 | 1.06 | 1.40 | 1.30 | 1.01 | | 1.25 | 0.87 | 1.54 | 1.34 |
| 12 | 1.38 | 1.45 | 1.40 | 1.82 | 1.19 | 1.19 | 1.43 | 1.03 | 1.41 | 1.31 | 1.25 | | 1.12 | 1.52 | 1.23 |
| 13 | 1.59 | 1.21 | 1.01 | 1.66 | 0.90 | 1.03 | 1.00 | 1.20 | 1.04 | 0.99 | 0.87 | 1.12 | | 1.37 | 1.01 |
| 14 | 1.28 | 1.09 | 1.34 | 1.31 | 1.24 | 1.51 | 1.15 | 1.60 | 1.36 | 1.26 | 1.54 | 1.52 | 1.37 | | 1.26 |
| 15 | 1.58 | 1.22 | 1.20 | 1.38 | 1.10 | 1.16 | 1.11 | 1.45 | 1.28 | 1.17 | 1.34 | 1.23 | 1.01 | 1.26 | |
| **Mean** | **1.55** | **1.30** | **1.27** | **1.63** | **1.17** | **1.27** | **1.19** | **1.43** | **1.34** | **1.19** | **1.29** | **1.34** | **1.14** | **1.34** | **1.25** |

**b**

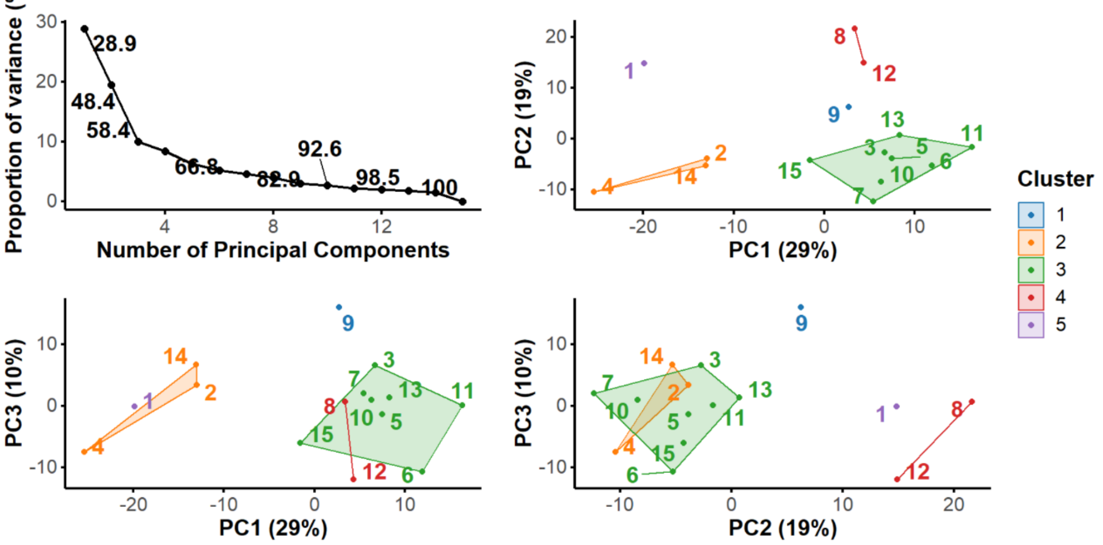


**Supplementary Fig. 6 | Comparative analysis of the 15 models in the ensemble.** (**a**) Pairwise RMSD for all heavy atoms for the final 15 structures of the dopamine-bound RKEC1 ensemble. (**b**) PCA analysis on the coordinates of the 15 starting conformers, used to identify two alternative starting conformations structurally as distant as possible from the initially selected conformer 1 (lowest-energy CNS/ARIA conformer) for the rMD simulations. The Scree plot (top left panel) shows the contribution of each principal component (PC) on the total variance and the cumulative variance. At least 10 PC are necessary to account for > 90% of the variance. For the sake of conciseness, the PCA coordinates of conformers were plot on two dimensions for the first three PCs only. However, resulting clusters determined by k-*means* (colored) and Euclidean distances between conformers were determined in a 10-dimensional space. Conformer 3 was found to be the furthest from 1, and conformer 9 to be the furthest from both 1 and 3.

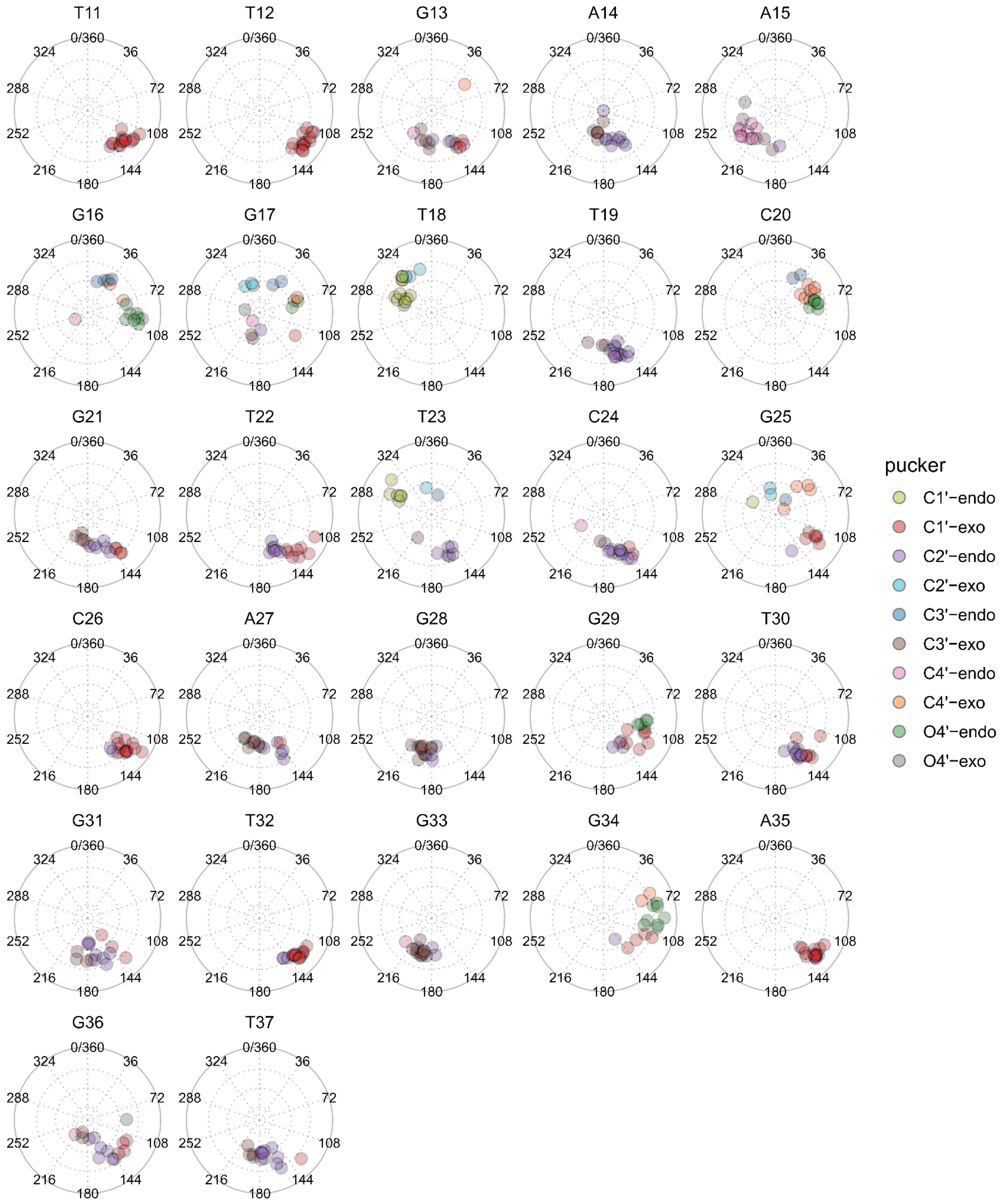


**Supplementary Fig. 7 | Sugar pucker analysis of the 15 minimized structures.** Sugar puckering calculated with the Altona & Sundaralingam method.(Altona & Sundaralingam, 1972) Distance from the center represents the amplitude of the sugar pucker, with planarity in the center.

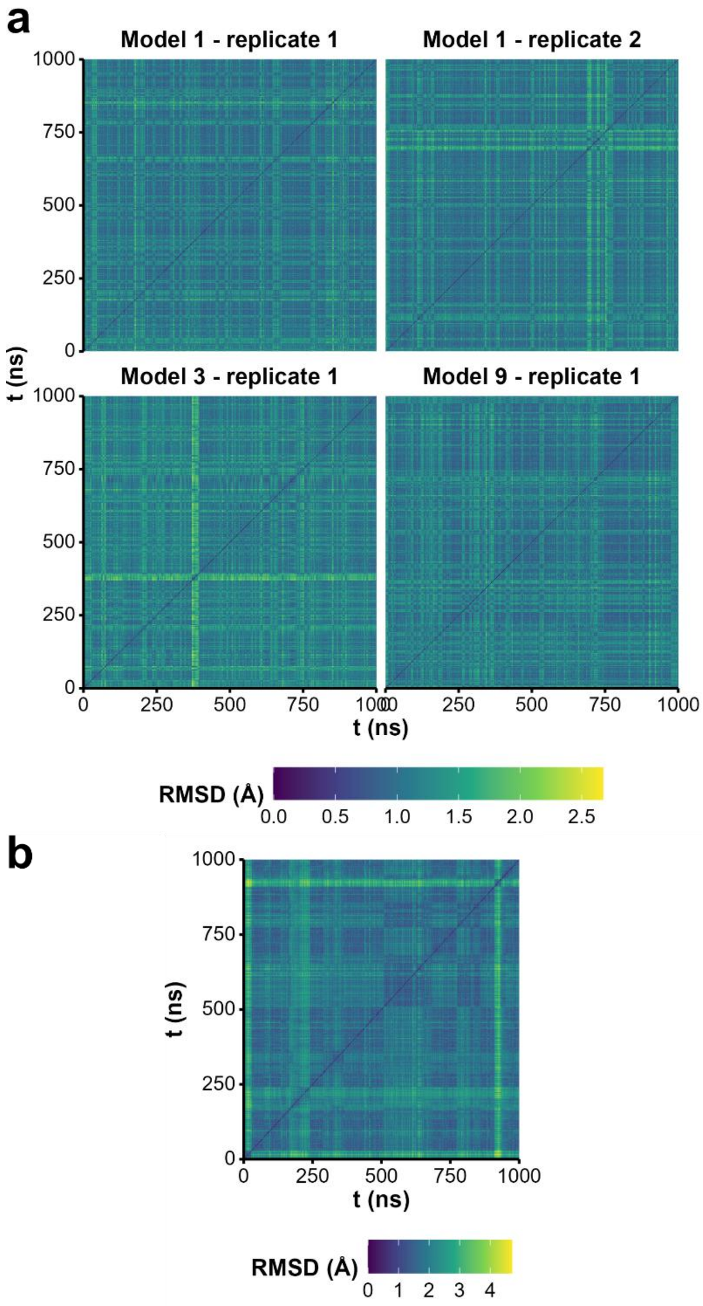


**Supplementary Fig. 8 | RMSD analysis during the MD simulations.** (**a**) RMSD calculated during the four restrained MD (rMD) microsecond simulations. Full pairwise heavy-atom RMSD matrices for the trajectories, shown as heatmaps. Square regions along the diagonal indicate intervals of structural similarity within each trajectory. (**b**) Similar RMSD analysis for the microsecond unrestrained MD (uMD) simulation.

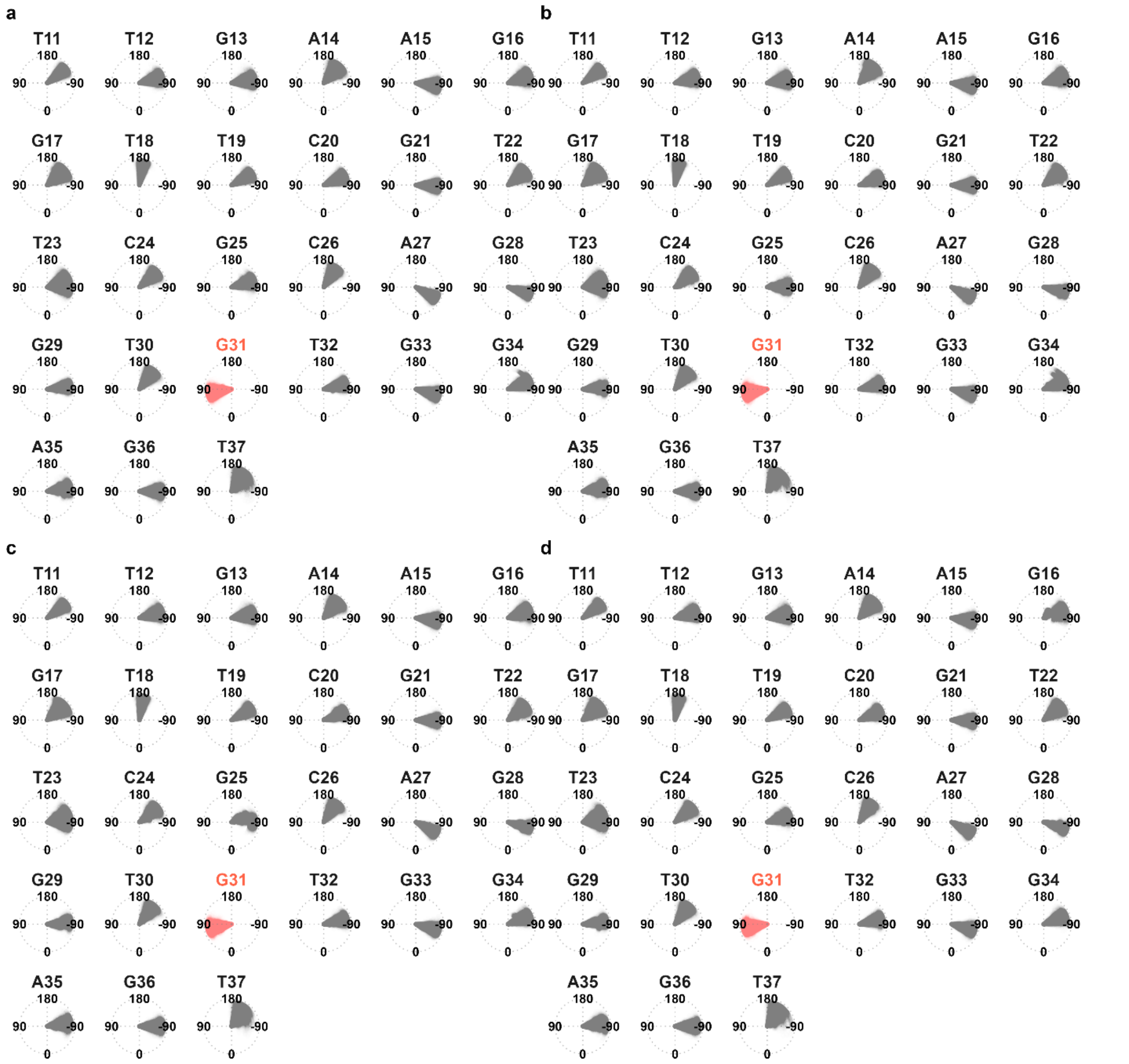


**Supplementary Fig. 9 | Chi dihedral angles of all residues during the four restrained MD microsecond simulations.** (**a**-**d**) Chi dihedral angles measured along the restrained MD (rMD) simulations performed with four replicates. The angles are indicated with a wrap in [-180;180], and proceed along the trajectory from center to outside of circle.

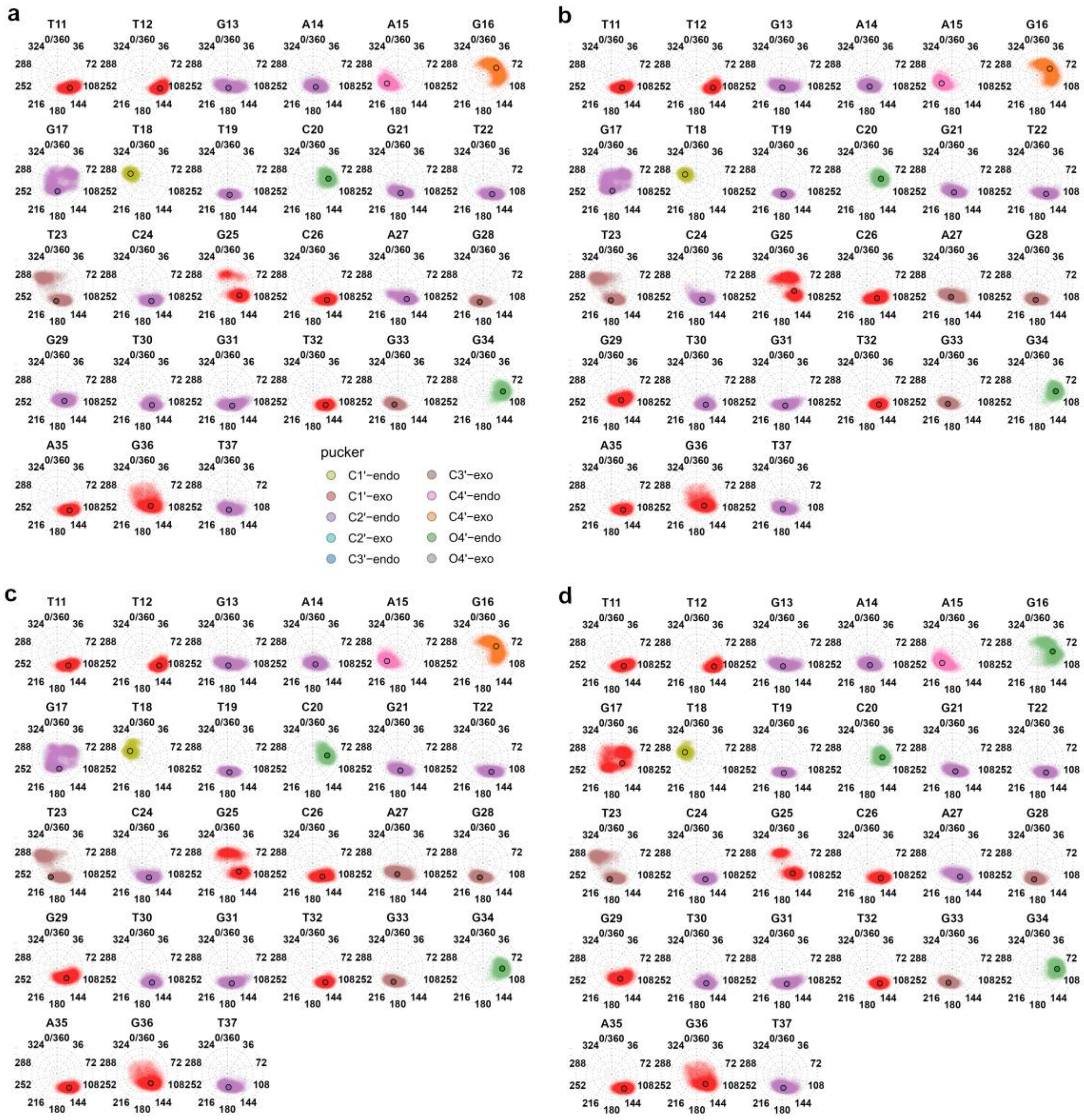

**Supplementary Fig. 10 | Sugar pucker values during the four restrained MD microsecond simulations. (a-d)** Sugar puckers calculated with the Altona & Sundaralingam method,(Altona & Sundaralingam, 1972) wrapped from 0 to 360 degrees, for the four restrained MD (rMD) simulation replicates. The position of the points relative to the center scales with the pucker amplitude (planarity in the center and largest amplitude at the border). The circled points represent the median pucker/amplitude. Calculation performed on 5000 evenly spaced frames.

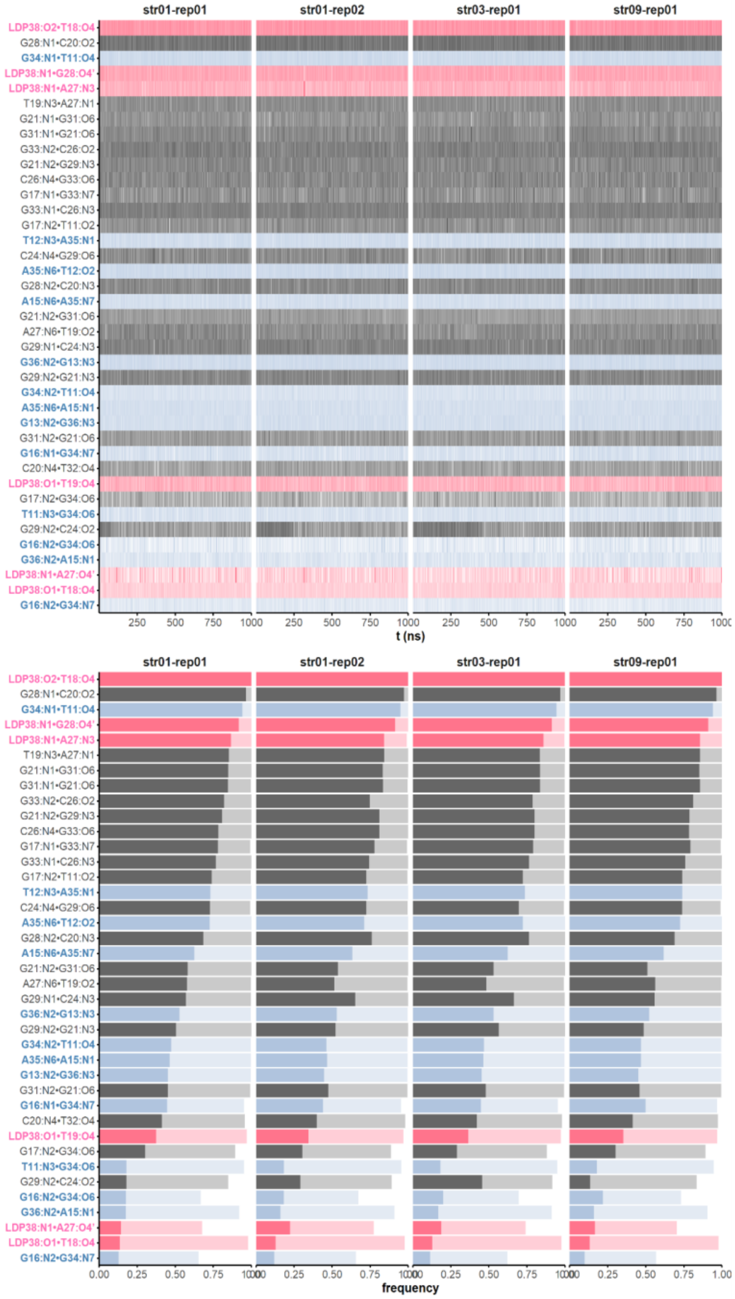


**Supplementary Fig. 11 | Formation of H-bonds over the course of the four restrained MD microsecond simulations**. The list of H-bonds was established on the minimized structures, with a maximum donor-acceptor distance of maximum 3.1 Å, keeping those formed in at least 20% of structures. H-bonds in the 5'- and 3'-end regions (residues 11:17 and 33:37) are colored in blue, and those involving the dopamine in pink. (**a**) Each line is a frame (one frame every 0.04 ns); the darker a frame, the shorter and more linear the H-bond (the transparency follows linearly the bond angle/length ratio). (**b**) Cumulative frequencies. The frequency of bond formation is shown for relaxed (3.5 Å and 120°; light colors) and more stringent (3.0 Å and 140°; dark colors) cut-offs. Only inter-residue H-bonds with a 'stringent' frequency above 0.10 are shown.

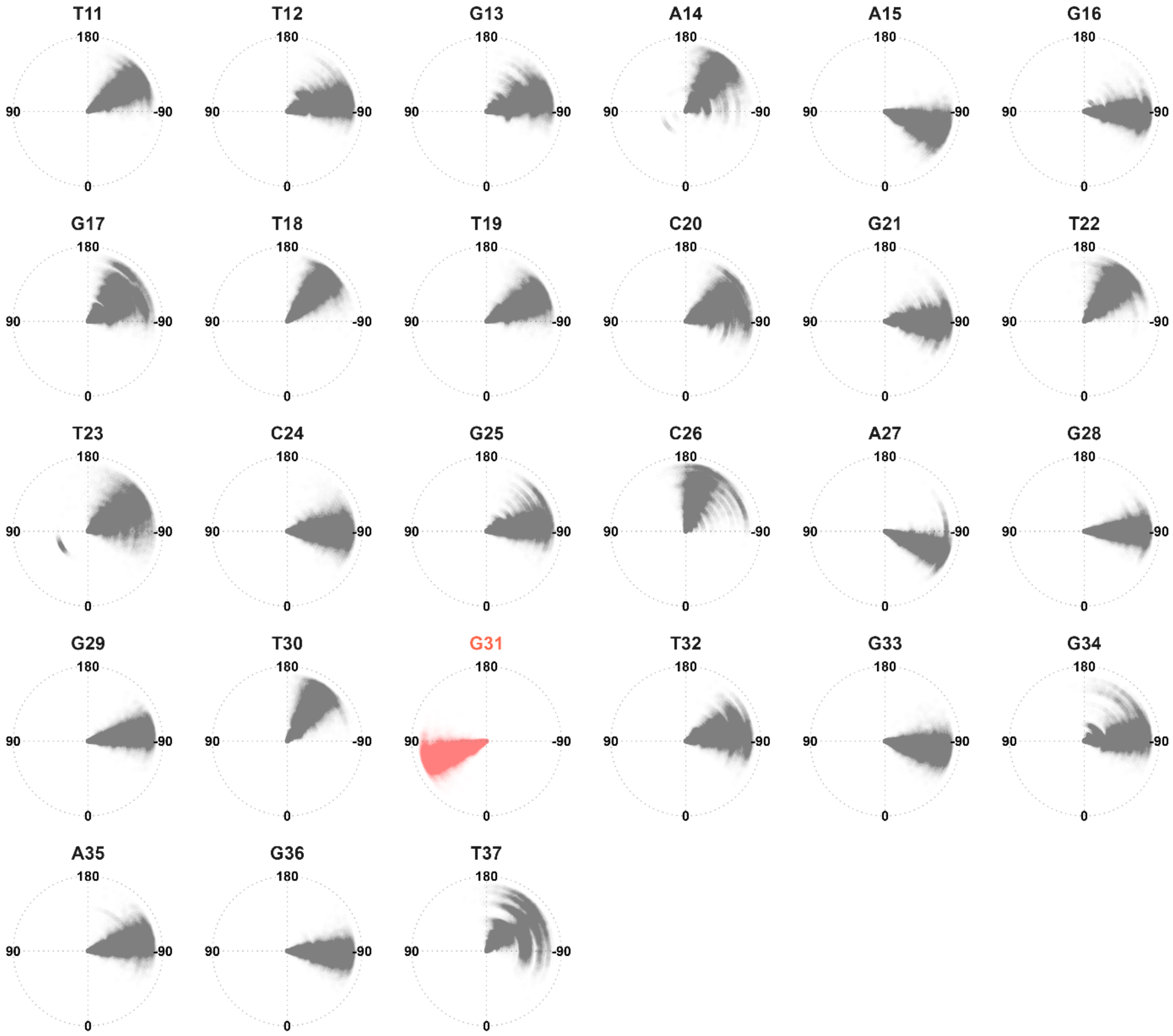


**Supplementary Fig. 12 | Chi dihedral angles of all residues during the microsecond unrestrained MD simulation.** Chi dihedral angles measured along the unrestrained MD (uMD) simulations performed with four replicates. The angles are indicated with a wrap in [-180;180], and proceed along the trajectory from center to outside of circle.

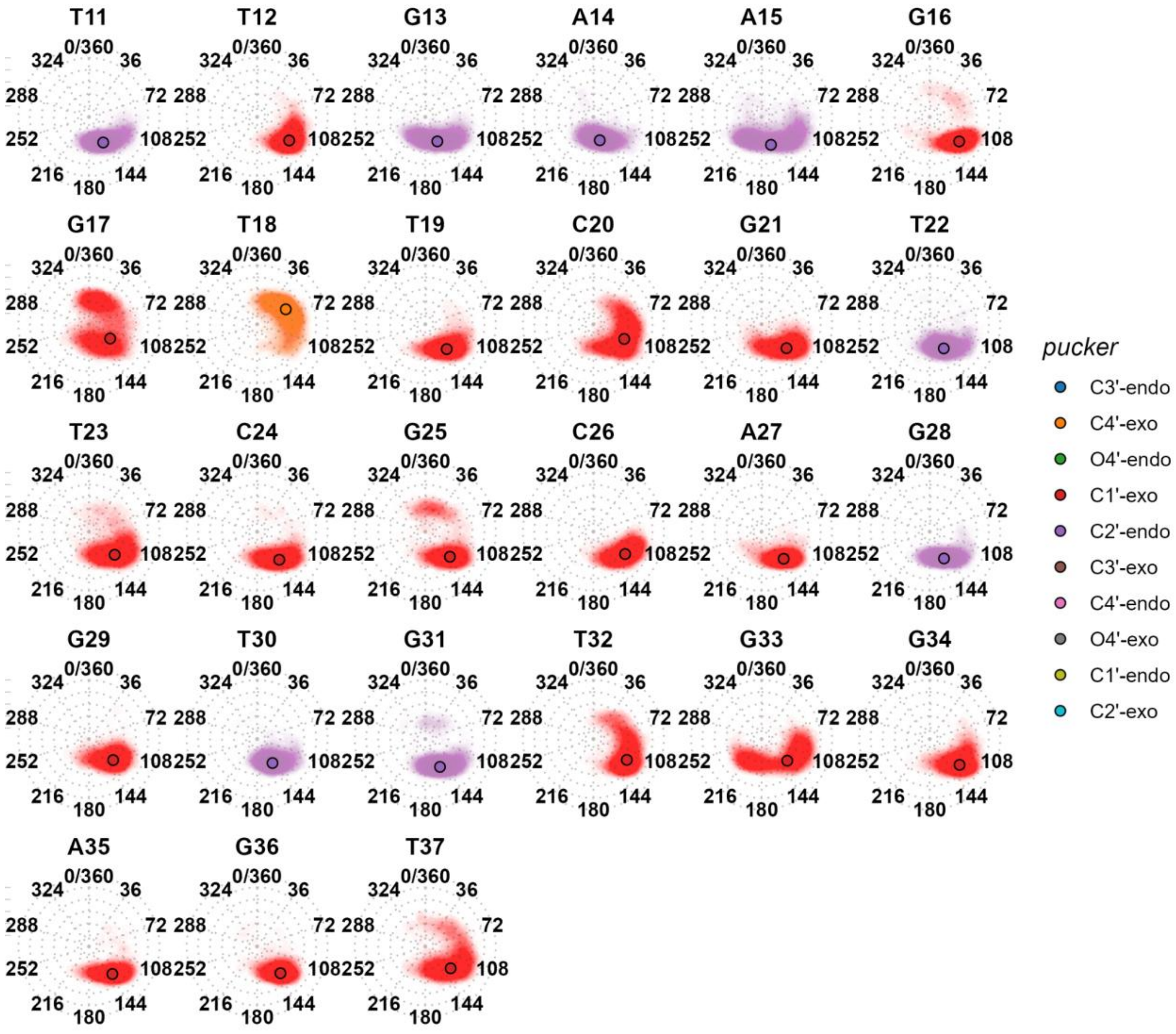


**Supplementary Fig. 13 | Sugar pucker values during the microsecond unrestrained MD simulation.** Sugar puckers calculated with the Altona & Sundaralingam method,(Altona & Sundaralingam, 1972) wrapped from 0 to 360 degrees, for the microsecond unrestrained MD (uMD) simulation. The position of the points relative to the center scales with the pucker amplitude (planarity in the center and largest amplitude at the border). The circled points represent the median pucker/amplitude. Calculation performed on 5000 evenly spaced frames.

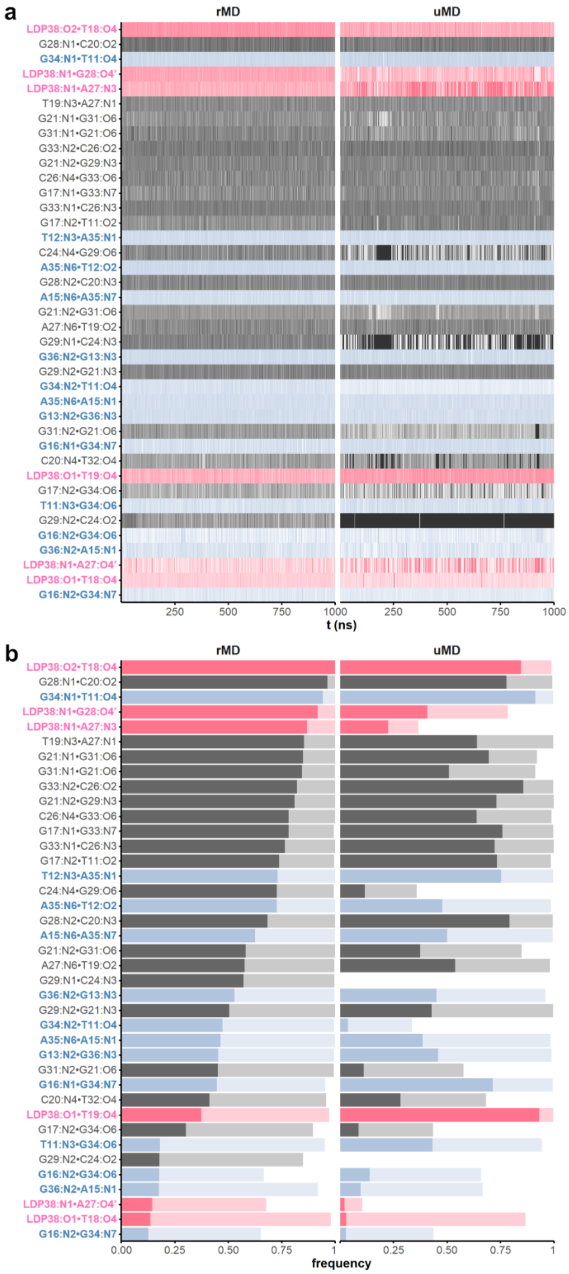


**Supplementary Fig. 14 | Formation of H-bonds over the microsecond unrestrained MD simulation**. The list of H-bonds was established on the minimized structures from the restrained MD (rMD) simulation (left column), with a maximum donor-acceptor distance of maximum 3.1 Å, keeping those formed in at least 20% of structures. H-bonds in the 5'- and 3'-end regions (residues 11:17 and 33:37) are colored in blue, and those involving the dopamine in pink. (**a**) Each line is a frame (one frame every 0.04 ns); the darker a frame, the shorter and more linear the H-bond (the transparency follows linearly the bond angle/length ratio). (**b**) Cumulative frequencies. The frequency of bond formation is shown for relaxed (3.5 Å and 120°; light colors) and more stringent (3.0 Å and 140°; dark colors) cut-offs. Only inter-residue H-bonds with a 'stringent' frequency above 0.10 are shown.

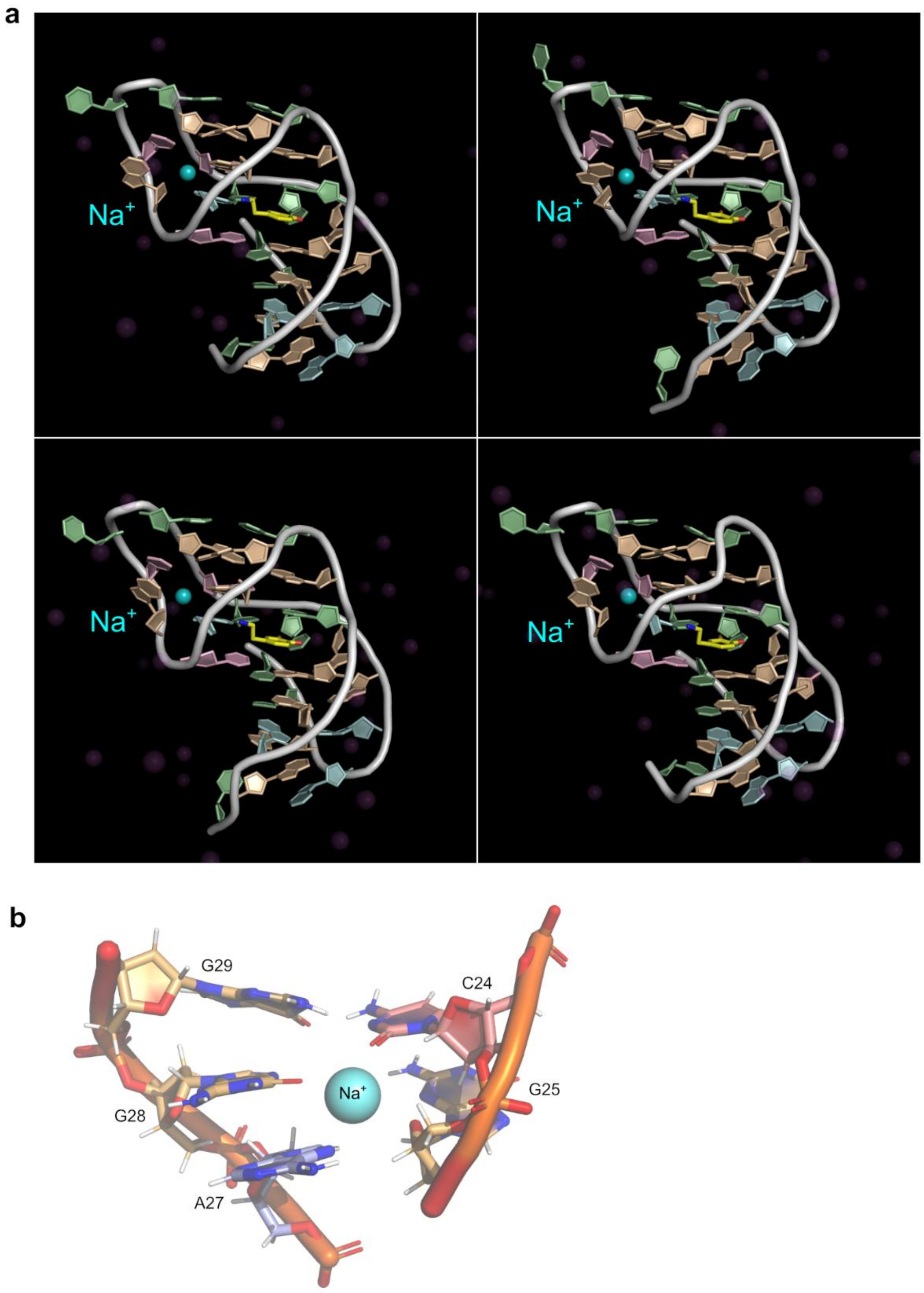


**Supplementary Fig. 15 | Observation of a stable sodium ion in the rMD simulations.** (**a**) A bound sodium cation can be observed at different times within all four restrained MD simulations. For example, the image captures frame 3486 from all four replicates. (**b**) The sodium cation contacts nucleotides in the loop region above the Watson–Crick C26–G33 base pair, and involves C24, G25, A27, G28 and G29. The rMD simulations were performed in the absence of magnesium. It is possible that this site could represent a bound magnesium ion in the NMR sample.

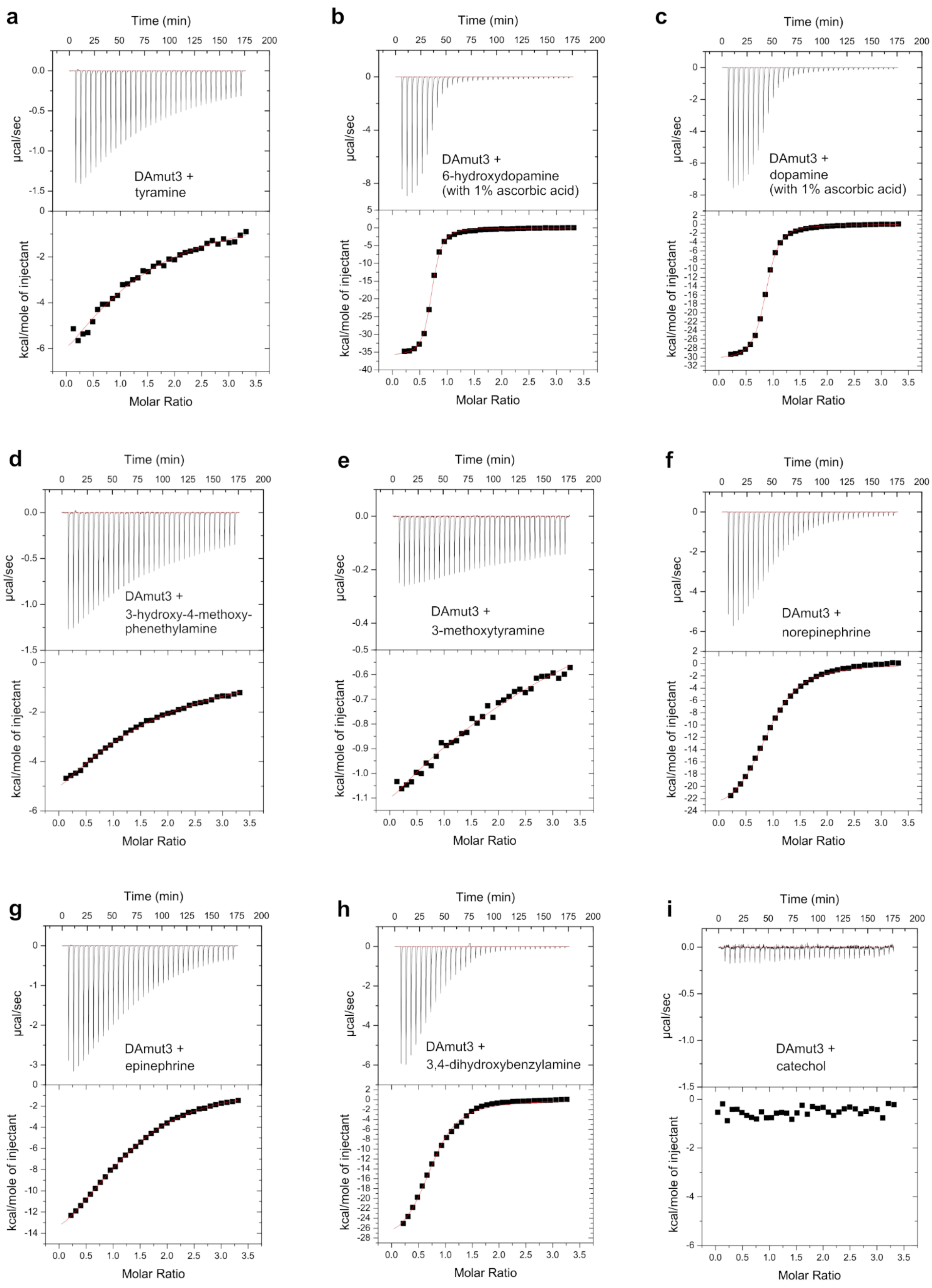


*(continued on next page)*

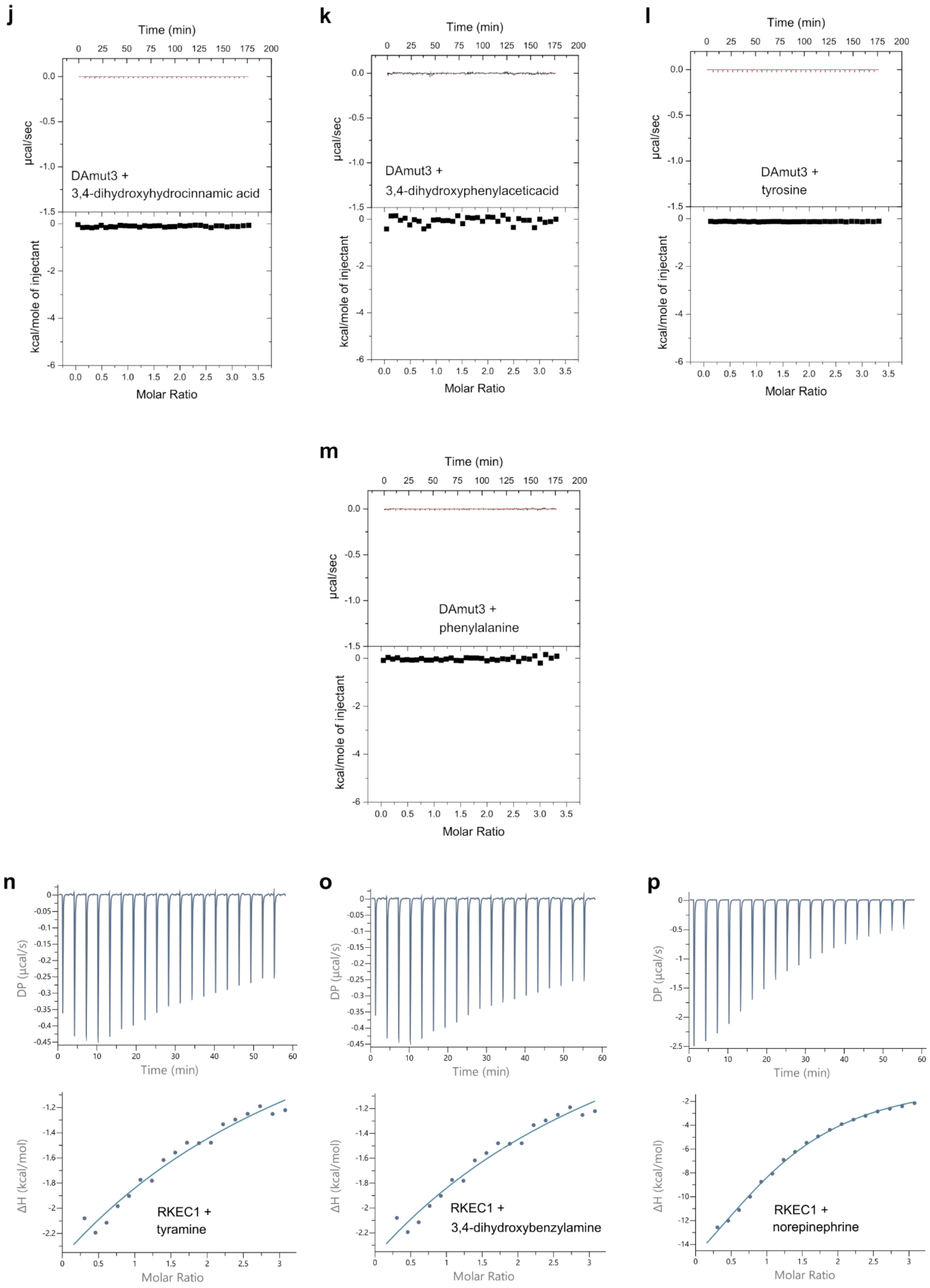


**Supplementary Fig. 16 | Representative ITC data related to Supplementary Table 1.**

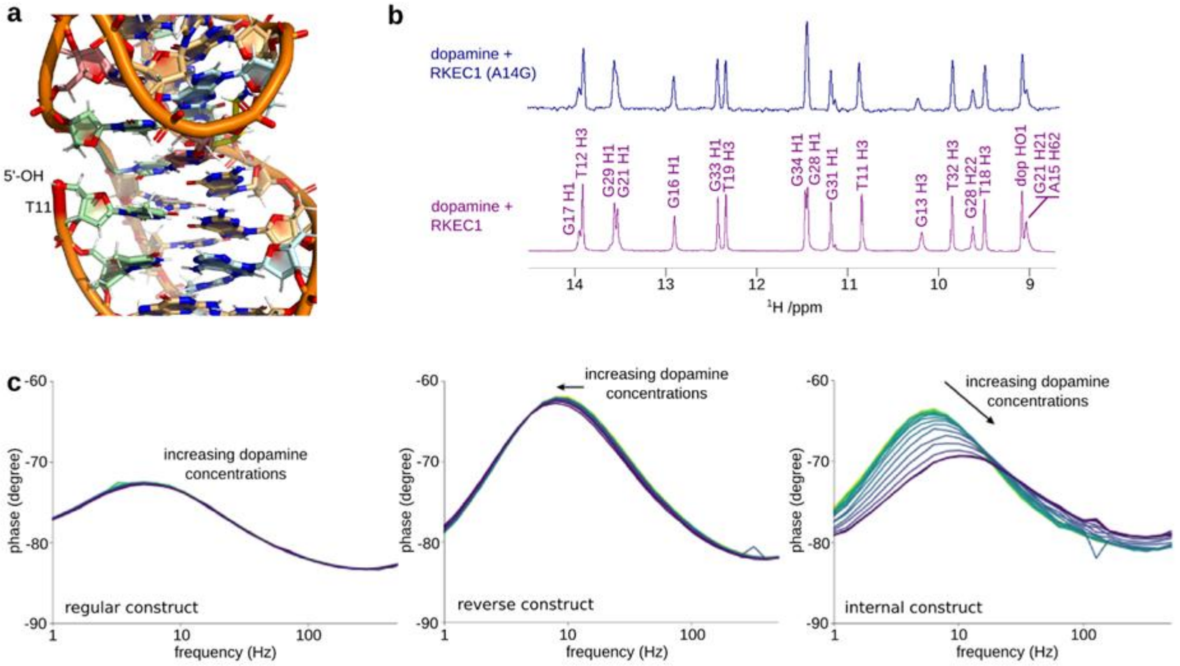


**Supplementary Fig. 17 | E-AB sensor design and phase-angle shift measurements.** (**a**) Position of the 5' hydroxyl in the dopamine-bound RKEC1. Note the proximity to the phosphate groups in the opposing strand. (**b**) In keeping with the lack of specific base interactions at position 14, the G14A RKEC1 variant maintains the ability to bind dopamine with an essentially unchanged spectrum in the imino region as compared to the wild-type RKEC1 complex. (**c**) Change in phase angle shift with increasing dopamine concentrations for the RKEC1-based E-AB biosensors with the MB regular, MB inverse, and MB internal arrangement. MB regular and MB inverse constructs displayed minimal phase angle shifts. In contrast, the MB internal construct displayed significant changes in the measured phase angle shift with increasing amounts of dopamine from either a change in the environment of the reporter or an alteration in its proximity with the electrode surface.
